# *DNMT3A*-R882 mutations intrinsically drive dysfunctional neutropoiesis from human haematopoietic stem cells

**DOI:** 10.1101/2025.10.08.679225

**Authors:** Giovanna Mantica, Aditi Vedi, Amos Tuval, Hector Huerga Encabo, Daniel Hayler, Aleksandra Krzywon, Emily Mitchell, William G Dunn, Tamir Biezuner, Hugo P Bastos, Kendig Sham, Antonella Santoro, Joe Lee, Nick Williams, Adi Danin, Noa Chapal, Yoni Moskovitz, Andrea Arruda, Edoardo Fiorillo, Valeria Orru, Michele Marongiu, Eoin McKinney, Francesco Cucca, Matthew Collin, Mark Minden, Peter Campbell, George S Vassiliou, Margarete A Fabre, Jyoti Nangalia, Dominique Bonnet, Liran Shlush, Elisa Laurenti

## Abstract

Clonal haematopoiesis (CH) arises from the expansion of hematopoietic stem cells (HSCs) carrying leukaemia-associated somatic mutations. CH is linked to pathological immune dysregulation and a greater risk of age-related inflammatory diseases. Yet, how CH mutations impact HSC differentiation into immune effector cells remains understudied. Here, we report a single-cell resolution functional and multi-omic investigation of HSC clonal and differentiation dynamics in individuals with *DNMT3A-R882* CH. *DNMT3A-*R882 reshapes the clonal architecture of haematopoiesis towards an aged phylogenetic structure. Functionally, *DNMT3A-*R882 HSCs produce decreased monocytic output but more abundant and mature neutrophil progeny compared to WT HSCs in the same individual. Whereas *DNMT3A-*R882 myeloid progenitors display attenuated inflammatory transcriptional programmes, *DNMT3A-* R882 mature neutrophils acquire proinflammatory and immunomodulatory features typical of maladaptive immunity and CH co-morbidities.

Our findings, validated in humanised mice, identify aberrant *DNMT3A*-R882 HSC-driven neutropoiesis as a key link between CH, immune dysregulation and risk of inflammatory disease.

## Introduction

The contribution of individual haematopoietic stem cells (HSCs) to blood production varies over the human lifespan. With ageing, somatic mutations are stochastically acquired, leading to the expansion of HSC clones carrying mutations conferring a competitive advantage. By the age of 70 years, haematopoiesis becomes oligoclonal, with around 30% of blood production arising from 20 or fewer mutant HSC clones^1^. When expanded clones carry a mutation in genes known to drive leukaemia and are present in more than 4% of blood cells (variant allele frequency, VAF ≥2%), this phenomenon is called clonal haematopoiesis of indeterminate potential (CHIP). Although blood counts remain normal or mildly perturbed, CHIP increases the relative risk of haematological malignancy by 10 fold, but the absolute risk is low (0.5-1% per annum cumulative risk), with a long latency period ^2,3^.

A large part of CHIP morbidity and mortality is attributed to an increased risk of non-haematological conditions, including cardiovascular disease^4^, chronic liver disease^5^, gout^6^, osteoporosis^7^, periodontitis^8^, chronic obstructive pulmonary disease^9^, and solid cancer ^10–12^. All these pathologies are underpinned by local and/or systemic chronic inflammation, indicative of immune dysregulation. Chronic inflammation is a hallmark of ageing ^13^ and is a known modulator of HSC function, leading to both clonal expansion of HSCs carrying CHIP mutations and acute and progressive loss of HSC regenerative capacity^14–18^. As in CHIP, HSCs from the mutant clone produce inflammatory cells, establishing how CHIP mutations affect the production and function of immune cells is essential for future prevention of leukaemia and CHIP comorbidities.

Mutations in the *DNMT3A* gene are the most common CHIP drivers ^2,3,19,20^ and are found in 18% of adult *de novo* AML^21^. Missense and truncating mutations occur across the length of the *DNMT3A* gene^22^, with substitutions at arginine 882 to histidine (R882H) or cysteine (R882C) being the most prevalent. R882 hotspot mutations decrease DNMT3A methyltransferase activity by 80% ^23^ and carry the highest risk of progression to AML ^2,24–26^. They also alter CpG flanking sequence recognition, redirecting residual methyltransferase activity to different sites than the WT^27,28^, and recruit PRC1 leading to reduced transcriptional activity and changed chromatin accessibility^29^.

HSCs carrying *DNMT3A* mutations are positively selected over the lifespan^1,30,31^. This increased fitness can be explained by *DNMT3A* loss of function conferring a self-renewal advantage to HSCs. Mouse HSCs in which *Dnmt3a* is deleted expand over serial transplantation, with multilineage differentiation progressively impaired in favour of self-renewal^32,33^. In mice with conditional activation of *Dnmt3a^R^*^878^*^H^*, the murine equivalent of *DNMT3A^R^*^882^, HSC gain a self-renewal advantage and differentiation is skewed towards myelopoiesis ^34,35^. Similarly, human primary HSCs carrying *DNMT3A* mutations display a competitive repopulation advantage upon transplantation^18,36,37^.

The subtle and progressive phenotypes associated with full or partial loss of function of *DNMT3A* are thought to arise from multiple mechanisms including progressive changes in methylation accrued slowly over time^38^, chromatin openness, and alternative splicing alterations ^29,39^. Indeed, deletion of *Dnmt3a* and *Dnmt3a/DNMT3A* mutations lead to relatively small effects at the transcriptome level^28,35,40–42^, and the alterations in the composition of HSC and progenitor (HSPC) compartment are also mild^28,40–43^. Whether the distinct composition of the HSPC pool, as well as lineage and transcription factor binding biases^28^ induced by *DNMT3A*-R882 result in functionally altered differentiation dynamics has not been studied to date in humans. Further, the clonal advantage of *Dnmt3A*-null or *Dnmt3A/DNMT3A* mutated HSCs is known to be exacerbated by inflammatory stimuli^15,18,44,45^, owing to the attenuated inflammatory responses observed in these cells compared to their WT counterparts^42^. Macrophages produced by *Dnmt3A/DNMT3A* mutated HSCs instead are thought to contribute a pro-inflammatory environment ^46–48^, which in turn may enhance the clonal advantage of mutated HSCs. It is likely that other *DNMT3A* mutant immune cell types contribute to this vicious circle, but the effect of *DNMT3A*-R882 on human HSCs differentiation towards innate immune cells has not been characterised.

Here, we studied how *DNMT3A*-R882 mutation alters the clonal and differentiation dynamics of human HSC/MPPs at single cell resolution. We identify perturbed myelopoiesis kinetics, driving production of innate immune cells with pro-inflammatory, myelosuppressive and immune modulating features, akin to those induced by maladaptive trained immunity.

## Results

### Clonal dynamics of DNMT3A*-R882H* mutant HSPCs

To assess how the acquisition of *DNMT3A*-*R882* mutation may affect HSC clonal dynamics over life, we performed phylogenetic analysis on HSPCs from a 63-year-old female with a large *DNMT3A*-R882H clone (VAF 34% in mobilised peripheral blood mononuclear cells). Following CD34+ selection, single HSCs/MPPs (Live/CD19-/CD33-/CD34+/CD45dim/CD38-/CD45RA-) and GMPs (Live/CD19-/CD33-/CD34+/CD45dim/CD38+/CD7-/CD10-) were isolated by FACS (**Fig.S1A**) and cultured *in vitro* for 3 weeks under conditions that permit simultaneous differentiation of erythroid, monocytic and granulocytic cells^49^ (**Fig.S1B**).

DNA isolated from 229 of the single-cell derived colonies was subjected to whole genome sequencing (WGS) with a depth of 13X, and for 96 of them, methylation status was assessed by EM-Seq^50^. 127 colonies passed QC for WGS and were used to reconstruct a phylogenetic tree using our previously published approach^1,51^. This tree displays an extensive *DNMT3A*-R882H clade (74% of all colonies), established very early in life, likely *in utero* (**Fig.1A**). *DNMT3A*-R882 mutations are known to lead to genome hypomethylation^23,28^. Consistently, colonies derived from single *DNMT3A R882* mutant HSC/MPPs or GMPs presented significantly lower levels of global methylation across all autosomes (**Fig.1B**). Within the *DNMT3A*-R882H clade, subclonal expansions of *TET2*-M1133K and *JAK2*-V617F were identified. Interestingly, the HSC/MPPs in the *DNMT3A*-R882H *JAK2*-V617F clade expressed significantly higher levels of CD49f, a marker of the most repopulating HSCs^52^, than *DNMT3A*-R882H or *DNMT3A*-WT HSC/MPPs (**Fig.S1C**). This is consistent with *JAK2-*V617F mutations preferentially expanding the most primitive HSCs^53,54^. In contrast, the *TET2-*M1133K clade was predominantly composed of GMP-derived colonies, in line with previous observations of *TET2* mutations driving clonal expansion preferentially in this progenitor compartment ^42^.

**Fig 1:**
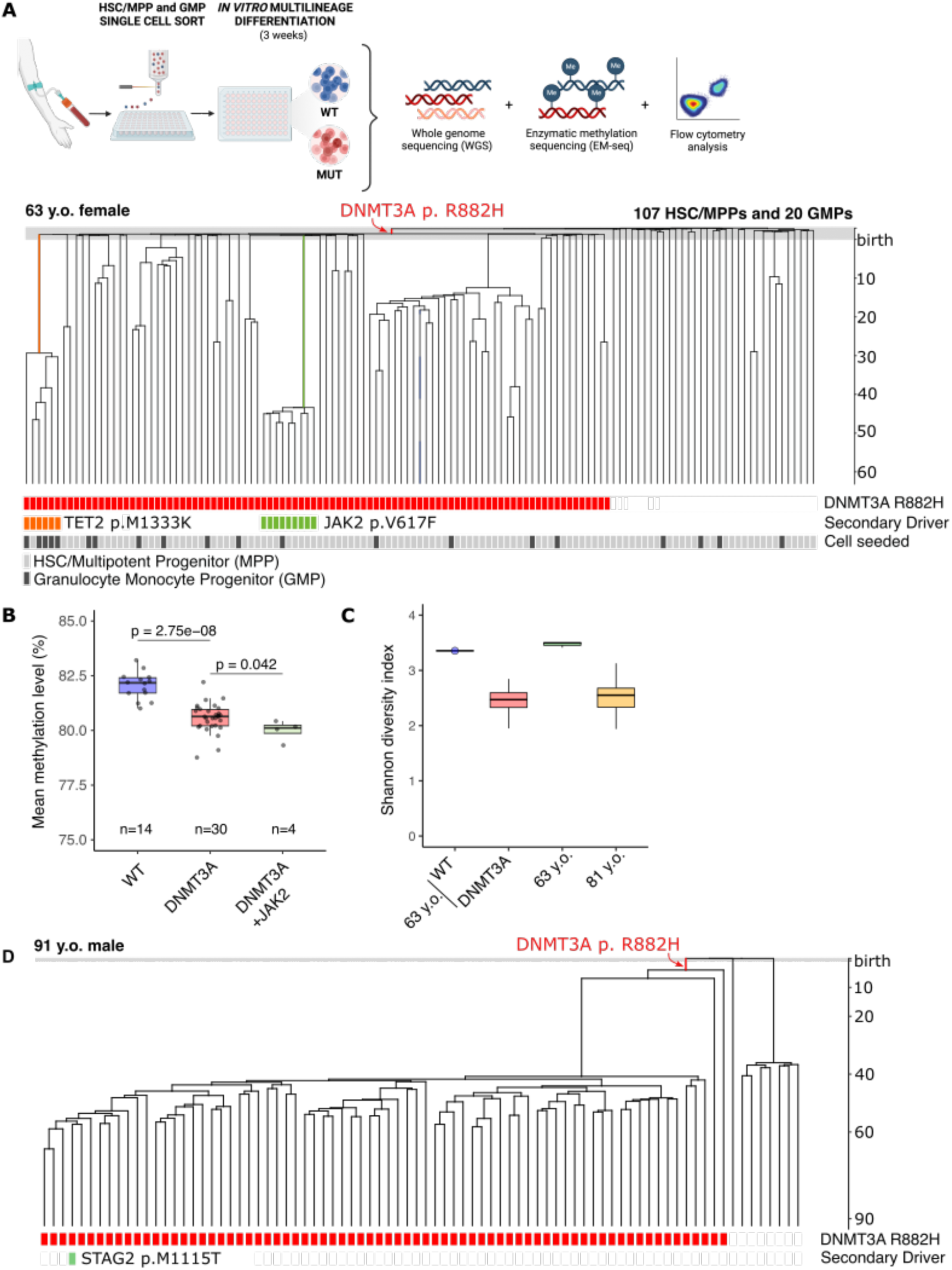
Clonal haematopoietic architecture and phylogenetic history of two individuals with *DNMT3A-* R882 CHIP. **(A)** Experimental design for single-cell characterisation of CHIP1 donor (top) and CHIP1 phylogenetic tree computed from 107 single HSC/MPP and 20 single GMP derived colonies (bottom). **(B)** Mean global methylation levels of CHIP1 myeloid colonies, stratified by genotype (autosomal DNA only; Wilcoxon rank-sum test; n = number of colonies). **C)** Shannon Diversity Index (SDI) of *DNMT3A*-R882 and WT sections of the CHIP1 tree (A), and from phylogenetic trees of two additional individuals aged 63 and 81yo from ^1^. SDI values for CHIP1 MUT section and full trees were calculated on 100 random subsamples matching the CHIP1 WT section size. **(D)** Phylogenetic tree of CHIP2, generated from 82 single-picked colonies grown for 14 days in semisolid media. In each tree **(A, D)**, each tip corresponds to a single colony; colonies sharing somatic mutations converge on a common ancestral branch, defining a clade. Y axes are scaled to time using the mutational rate as a molecular clock, as in ^1^. Bars under the trees show colonies mutant for *DNMT3A* and for secondary myeloid drivers. In **(A),** additional information regarding the cell of origin of the colony is provided in the bottom bar: HSC/MPPs (grey) or GMPs (dark grey).

Interestingly, the *DNMT3A*-R882H mutant clone exhibits a complex clonal architecture with multiple subclonal expansions, absent in the WT compartment. The Shannon Index, a measure of clonality, differed significantly for the *DNMT3A*-R882H and WT portions of the tree. Whereas the Shannon index of the WT portion of the tree was similar to that of a 63-year-old healthy individual^1^, the Shannon index for the *DNMT3A*-R882H clade was significantly lower, and comparable to that of an 81 year old^1^ (**Fig.1C**). This indicates that acquisition of *DNMT3A*-R882H mutation in HSCs leads to premature oligoclonality, possibly favouring expansion of additional mutations. We then performed phylogenetic analysis from peripheral blood of a 91-year-old male, who had been serially monitored in the context of the SardiNIA longitudinal study^55^, and was found to have progressive expansion of the *DNMT3A*-R882H clone from 74 to 89 years of age (CHIP2, VAF expanding from 5% in mononuclear cells at 74 years of age to 34% at 89 years of age)^30^. Again, we observed acquisition of *DNMT3A*-R882H within the first 5 years of life. The *DNMT3A*-R882H clone had further expanded over two years (45% VAF at 91 years old), with one prominent subclonal expansion emerging from the *DNMT3A*-R882H clade later in life (**Fig 1D**). Interestingly, while it was not possible to perform longitudinal sampling on CHIP1, four years post-sampling this donor developed essential thrombocythemia originating from the JAK2 V617F clonal expansion successive to *DNMT3A-R882*. The pattern of clonal evolution in both donors suggests clonal sweeps occurring due to secondary mutations acquired decades after *DNMT3A-R882* acquisition.

### DNMT3A*-R882* mutant HSC/MPP proliferation and commitment towards the erythroid and myeloid lineages are similar to WT HSC/MPPs

As we recorded cell surface phenotyping data from the 107 HSC/MPP derived colonies used to generate the phylodynamic tree of CHIP1, we next investigated whether the expression of specific differentiation markers in the differentiated colonies associated with specific HSC/MPP genotypes. Whereas no significant differences were found for GlyA or CD15 median fluorescence intensity (MFI; not shown), the MFI of the monocytic marker CD14 was significantly lower in myeloid colonies (CD45+/CD56-) derived from *DNMT3A*-R882H HSC/MPPs than in those derived from *DNMT3A*-WT HSC/MPPs (**Fig.2A**). These data suggest that *DNMT3A*-R882H HSC/MPPs may differ from WT HSCs in their myeloid differentiation dynamics.

**Fig 2:**
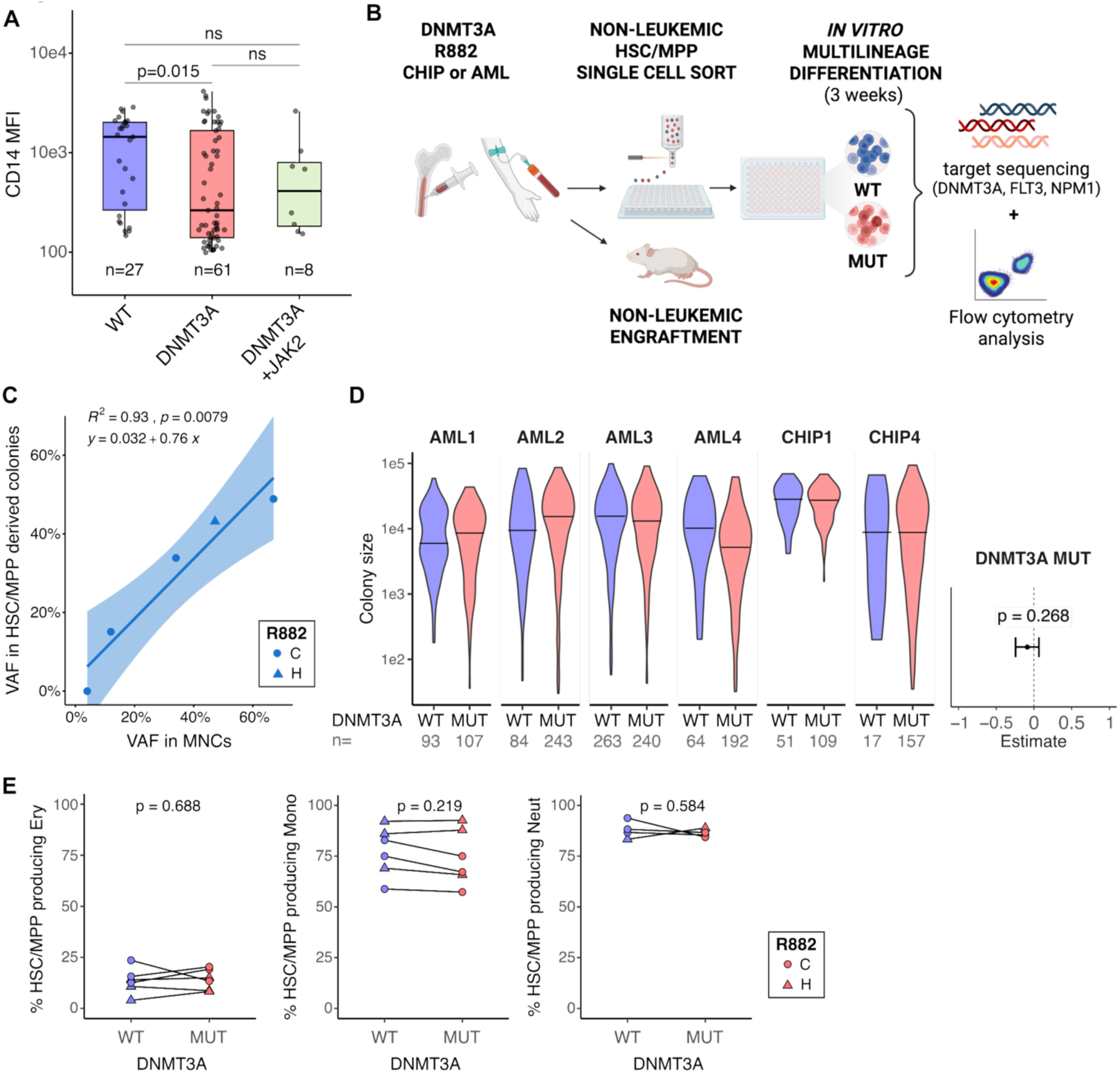
*DNMT3A-R882* mutations do not confer selection or differentiation biases to HSC/MPPs. **(A)** CD14 mean fluorescence intensity (MFI) within CD45+/GlyA-/CD56-cells of CHIP1 HSC/MPP derived colonies from the indicated genotypes (as determined from WGS, **Fig.1A**). p: two-sided t-test **(B)** Experimental design using primary samples from individuals carrying *DNMT3A-*R882 mutations to evaluate the effects of the mutation on HSC/MPP function. **(C)** *DNMT3A-*R882 variant allele frequency (VAF) measured in HSC/MPP derived colonies vs MNCs from PB of the same individuals with CHIP (n=5). p: R^2^ statistic representing fit with linear regression model. Equation of the regression line shown, with slope not significantly different from 1 (p=0.0079). **(D)** Colony size, measured as number of single cells, produced by single *DNMT3A-*R882 and WT HSC/MPPs for each individual. n= number of colonies analysed per condition. Forest plot report estimated *DNMT3A* mutation effect (with 95% confidence interval) on the parameter of interest, obtained by mixed effect modelling (see Methods). p values computed with Satterthwaite’s method. **(E)** Percentage of *DNMT3A-*R882 MUT or WT HSC/MPPs producing colonies containing erythrocytes (Ery), monocytes (Mono), and neutrophils (Neut) in each individual. n=6 (Ery, Mono), n=4 (Neut), p: paired two-tailed Wilcoxon signed-rank test.

To further explore differences in HSC/MPP function conferred by *DNMT3A*-R882 mutations, we established a cohort of 10 independent carriers of *DNMT3A*-R882 (6 R882H, 4 R882C; 5 CHIP, 5 AML; age range: 38-81 years, median age: 63 years; VAF range: 4-67%, median VAF: 45.9%; **Table S1**). In this cohort, we included AML samples from which we could confidently isolate pre-leukemic HSC/MPPs (preL-HSCs) carrying only the *DNMT3A*-R882 mutation, while excluding leukemic cells of any maturity. This was achieved by: i) selecting AML samples with blasts of CD34-phenotype; ii) selecting samples which we validated gave rise to multilineage engraftment in immunocompromised mice, indicative of high levels of preL-HSCs; iii) isolating CD45dim/CD34+/CD38-/CD45RA-HSC/MPPs, a published strategy for preL-HSCs^36^; iv) using culture conditions^49^ that do not support growth of leukemic stem cells; v) verifying by targeted sequencing that none of the single cell derived colonies carried *NPM1*, *FLT3*-ITD and other leukaemia-associated mutations present in the AML donors (**Table S1**).

Single HSC/MPPs were FACS sorted from CD34+ cells (**Fig.S1A**) and cultured *in vitro* for 3 weeks before phenotyping (**Fig.S1B**) and genotyping by targeted sequencing (**Fig.2A**). Overall, 5,671 single HSPCs were sorted from 10 samples, yielding 2,526 single cell derived colonies (**Table S2**). Colony genotypes were assigned after mutation calling using 3 independent variant callers (VarScan, Platypus and Mutect, **Table S2**). Median clonogenic efficiency ranged from 18-78% across all samples (**Fig.S2A**). We observed no differences in cell surface marker expression of HSCs/MPPs that did or did not produce colonies *in vitro* amongst all samples (**Fig.S2B**). Moreover, *DNMT3A*-R882 VAF calculated on the single cell derived colonies did not differ from the one of MNCs in CHIP individuals (**Fig 2C**). These two observations suggest that the culture conditions used do not select for a specific HSPC subset or mutational status.

Then we assessed the commitment and differentiation capacity of *DNMT3A*-R882 mutant HSC/MPPs across different lineages by analysing flow cytometry data from those samples with sufficient intra-individual statistical power (≥20 colonies per genotype). 1,620 colonies were included, obtained from 6 independent samples (2 CHIP and 4 AML, 3 *DNMT3A*-R882H and 3 *DNMT3A*-R882C, 38-78 years, 20-48% *DNMT3A*-R882VAF in MNCs, **Table S2**). At the time of index sorting, *DNMT3A*-R882 HSC/MPPs displayed slightly, but significantly, higher levels of CD49f and lower levels of CD34 cell surface expression, than their WT counterparts (**Fig.S2C-D**). Second, colony size, measured by flow cytometry, was similar between WT and *DNMT3A*-R882 mutant HSC/MPP derived colonies (**Fig.2D**) across all CHIP and AML individuals. This indicates that *DNMT3A*-R882 mutation does not affect the overall proliferation capacity of HSC/MPPs.

Mouse *Dnmt3A* knock-out^40^ and human *DNMT3A*-*R882* HSPCs^28^ have been shown to be transcriptionally biased towards erythropoiesis. Whether this translates into more HSC/MPPs committing towards the erythroid lineage at the expense of the myeloid has not been studied. We found similar proportions of *DNMT3A*-R882 mutant and WT HSCs/MPPs producing colonies containing erythroid cells (**Fig.2E**). Similarly, *DNMT3A*-R882 mutant and WT HSCs/MPPs produced similar proportions of colonies containing monocytes or neutrophils (**Fig.2E**). This indicates that, in our assay, HSC/MPPs with *DNMT3A*-R882 mutation can commit to the erythroid, monocyte and neutrophil lineages to the same capacity as WT HSC/MPPs.

### *DNMT3A*-R882 mutation enhances neutrophil differentiation of HSC/MPPs

Next, we investigated whether *DNMT3A*-R882 mutations quantitatively alter the output of mature cells produced by HSC/MPPs. First, from each individual donor we generated an *in silico* pool of all mature cells produced by single HSC/MPPs, akin to a landscape of differentiation, using CytoTree^56^. Dimensionality reduction was performed and a 2D UMAP embedding based on the expression of all cell surface markers was generated. WT and *DNMT3A*-R882 cells were found in all areas of these embeddings. Nonetheless, in some areas of the map, mature cells produced by *DNMT3A*-R882 HSC/MPPs were over-or underrepresented. High *DNMT3A*-R882:WT ratios largely corresponded to areas with high CD66b, a marker of neutrophils, while low *DNMT3A*-R882:WT ratio to areas with high CD14, a monocyte marker (**Fig.3A, Fig.S3A**).

**Fig 3:**
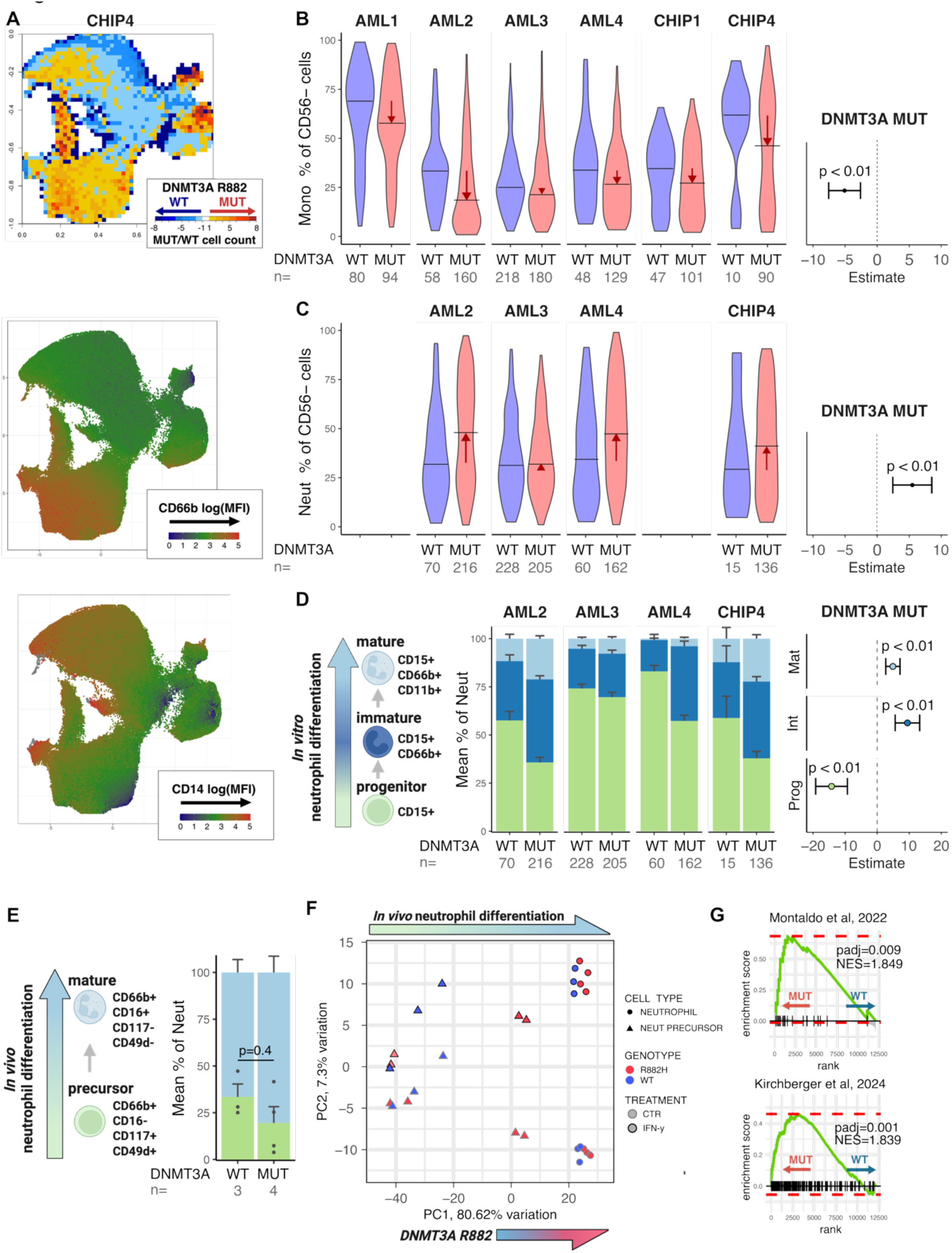
*DNMT3A-*R882 mutations affect HSC/MPP myeloid differentiation dynamics. **(A)** Representative example of pooled *in silico* analysis of all mature cells derived from HSC/MPPs from one individual (CHIP4, n= 287,681 cells). Top panel: ratio of *DNMT3A-*R882 mutated cells / WT cells in each area of the UMAP; middle and bottom: MFI of CD66b (middle) and CD14 (bottom) cell surface markers. **(B)** Percentage of monocytes, defined as CD14+/CD11b+ relative to myeloid cells (GlyA-/CD56-, **Fig. S1B**) within monocyte-containing colonies. **(C)** Percentage of neutrophils, defined as the sum of early (CD14-/CD15+/CD66b-), intermediate (CD14-/CD15+/CD66b+/CD11b-), and mature (CD14-/CD15+/CD66b+/CD11b+) neutrophils, relative to myeloid cells (GlyA-/CD56-) within neutrophil-containing colonies. **(B-C)** n = number of monocyte-containing (B) or neutrophil-containing (C) colonies per condition per individual, horizontal marks in the violin plot indicate medians; red arrows indicate the direction of effect of the mutation. **(D)** Composition of the neutrophil population in WT and *DNMT3A-*R882 MUT neutrophil-containing colonies. Mean and standard error of the percentage of early, intermediate and mature neutrophils (as defined in left panel). n= number of neutrophil-containing colonies analysed per condition per individual. In (**B-D),** forest plots report estimated *DNMT3A* mutation effect (with 95% confidence interval) on the parameter of interest, obtained by mixed effect modelling (see Methods). p values were computed with Satterthwaite’s method and corrected with Benjamini–Hochberg’s. **(E)** Relative composition of precursors and mature neutrophils in BM of mice transplanted with *DNMT3A*-R882 CRISPR edited HSCs and WT controls. p: Wilcoxon rank sum test. **(F)** Principal Component Analysis of bulk RNAseq data of precursors and mature neutrophils from (**E)**. **(G)** GSEA enrichment of neutrophil maturation signatures ^60,61^ in *DNMT3A*-R882 vs WT neutrophils from (**E)**.

We next analysed single cell derived colonies containing myeloid cells (**Fig.S1b**; see Methods for definition criteria) in each donor, employing conventional gating. The percentage of monocytes out of all myeloid cells, was overall about 5% lower in *DNMT3A*-R882 mutant HSC/MPP derived colonies compared to WT (95% CI: –7.55 to –2.60, p value<0.001, **Fig.3B**). In contrast, the percentage of neutrophils was approximately 5% higher in *DNMT3A*-R882mutant colonies than in WT colonies (95% CI: 2.76 to 8.82, p value<0.001, **Fig.3C**).

In addition to increasing relative neutrophil production, *DNMT3A*-R882 mutations may also influence neutrophil maturation. We quantified the production of three stages of neutrophil differentiation in the assay above (**Fig.S1b**, bottom panel): progenitors (GlyA-CD45+ CD56-CD14-CD15+ CD66b-CD11b-), intermediate (GlyA-CD45+ CD56-CD14-CD15+ CD66b+ CD11b-) and mature neutrophils (GlyA-CD45+ CD56-CD14-CD15+ CD66b+ CD11b+)^57,58^. Neutrophil composition was significantly shifted towards more mature subsets in neutrophil colonies derived from *DNMT3A*-R882 HSC/MPPs, whereas early neutrophils were more abundant in neutrophil colonies derived from WT HSC/MPPs (**Fig.3D**). Mutations in *TET2*, another gene strongly associated to CHIP, lead to the increased production of neutrophils with low granule complexity ^59^. However, in our dataset, neutrophil granularity was overall similar in WT and *DNMT3A*-R882 mutant neutrophils (**Fig.S3B**).

To validate these findings in an independent model, we introduced *DNMT3A*-R882 mutations in umbilical cord blood Lin-CD34+ CD38-HSPCs using CRISPR-Cas9 editing, then transplanted these cells in non-conditioned NSG-SGM3-cKit^W41/W41^ mice (**Fig.S3C**). Upon successful reconstitution of human haematopoiesis (**Fig.S3D)**, the degree of editing in individual mice was evaluated in BM human CD45+ cells (**Fig.S3D**). Neutrophil precursors (PRE, CD66b+ CD16-CD117+ CD49d+) and mature neutrophils (NEU, CD66b+ CD16+ CD15+) were then quantified. In accordance with the *in vitro* experiments performed with HSC/MPPs natively carrying *DNMT3A*-R882 mutations, we noted a trend towards more phenotypically defined mature neutrophils in *DNMT3A*-R882-edited animals (**Fig.3E**). Neutrophil precursors and neutrophils were also flow sorted and cultured overnight in the presence or absence of IFNγ, before performing bulk RNA-seq (**Fig.S3C, Fig.S3D**). Principal component (PC) analysis showed that PC1 represents maturation, with neutrophil precursors clustered away from neutrophils along this PC, while PC2 separated IFNγ-treated and untreated samples. Interestingly, *DNMT3A*-R882*-*edited neutrophils displayed mildly but significantly higher PC1 values than control neutrophils (**Fig.S3E**), suggesting increased maturity. Consistently, published neutrophil maturation transcriptional signatures^60,61^ were significantly enriched in *DNMT3A*-R882-edited neutrophils compared to control neutrophils (**Fig.3G**).

These data collectively show that, in two independent experimental systems, *DNMT3A*-R882 mutant HSC/MPPs produce relatively more neutrophils, which are also phenotypically more mature than their WT counterparts.

### Inflammatory transcriptional programmes are dampened in *DNMT3A*-R882 neutrophil progenitors but accentuated in stimulated mature neutrophils

To further characterise whether *DNMT3A*-R882 mutation in HSC/MPPs leads to the production of mature myeloid cells with distinct transcriptional properties and response to inflammation, colonies were generated from single HSC/MPPs from a CHIP donor (CHIP5) in the same conditions as above. After three weeks of culture, colonies were split, with half left untreated and the other treated with IFNγ overnight. IFNγ was chosen as exposure to it increases the clonal expansion of *Dnmt3A*-null HSCs^15^ and it is a potent activator of pro-inflammatory responses in neutrophils. Treated or untreated colonies of the same genotype were then pooled, myeloid populations were flow-sorted and submitted for 10X Genomics scRNA-seq (see methods for protocol modifications to ensure neutrophil capture)(**Fig. 4A**). 43,501 single cells passed quality control and were annotated using label transfer from Kwok et al. ^62^ (**Fig.S4A-C**, **Table S3)**. We could identify monocytes as well as 4 stages of neutrophil maturation, ranging from neutrophil progenitors (NP) to mature neutrophils (**Fig.4B and Fig.S4B**). As expected, *DNMT3A*-R882 mutant and WT cells were found across the whole UMAP (**Fig.S4A**). In addition, IFNγ treatment induced upregulation of IFN-responsive and inflammatory signatures across all myeloid subsets, as well proliferation signatures in NPs (**Fig.S4D**).

**Fig. 4:**
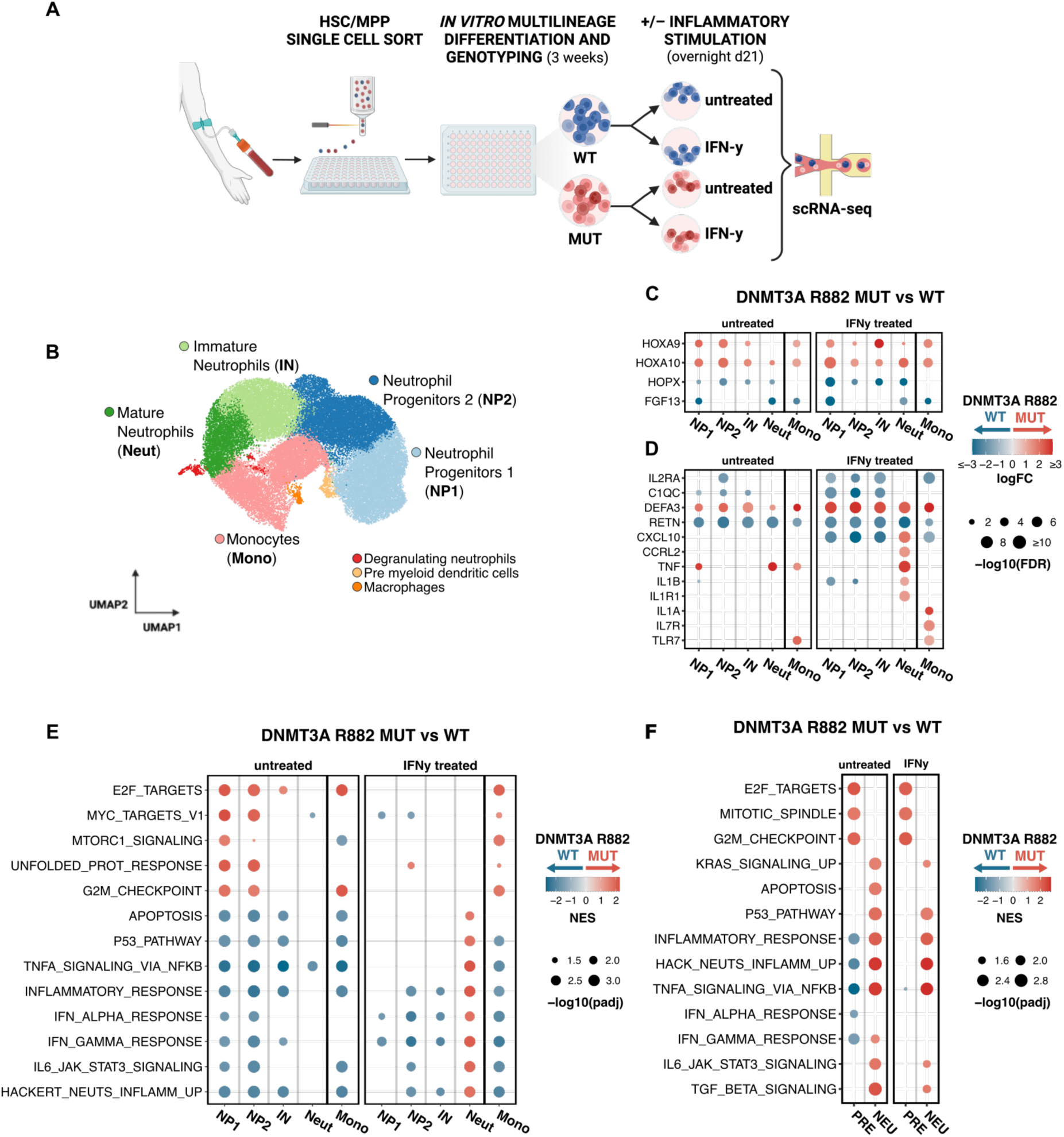
Inflammatory pathways are altered in *DNMT3A-* R882 monocytes and neutrophils at steady-state and upon inflammatory stimulus. **(A)** Schematic of the experimental design. HSC/MPPs were isolated from PB and differentiated *in vitro* for 3 weeks into single-cell derived colonies. Each colony was genotyped for the presence or absence of the *DNMT3A*-R882 mutation. At the end of the culture period, colonies were split, and half underwent overnight stimulation with IFNγ, prior to 10X chromium scRNA sequencing. **(B)** UMAP from scRNA-seq of 43,501 myeloid cells derived from *DNMT3A-*R882 or WT HSC/MPPs, treated or not treated with IFNγ, cells coloured by annotated cluster. **(C-D)** Dot plot of the expression of selected genes in the indicated clusters. All genes shown were differentially expressed between *DNMT3A-*R882 and WT untreated cells of the indicated clusters (FDR<0.05 and logFC >|1|). **(E)** GSEA analysis comparing *DNMT3A*-R882 vs WT cells in untreated (left) and IFNγ-treated (right) conditions within the indicated clusters. Positive NES scores (red) indicate enrichment in *DNMT3A-*R882 cells. NP1: Neutrophil Progenitors 1, NP2: Neutrophil Progenitors 2, IN: Immature Neutrophils, Neuts: mature Neutrophils; Mono: monocytes **(F)** GSEA analysis comparing precursors (PRE) and mature neutrophils (NEU) derived *in* vitro from CRISPR-edited *DNMT3A*-R882 HSCs vs WT in untreated (left) and IFNγ-treated (right) conditions.

We then performed differential gene expression analysis between *DNMT3A*-R882 mutant and WT cells in each of the myeloid clusters **(Table S3)**. The most marked differences in transcription were found in monocytes (with respectively >430 and >720 genes differentially expressed in *DNMT3A*-R882 untreated and IFNγ-treated monocytes compared to WT, FDR<0.05, |logFC|>1 **Fig.S4E**). Notably, despite the overall lower transcriptional activity of neutrophils, *DNMT3A*-R882-driven differences were also observed in this compartment (>80 and >280 genes differentially expressed in untreated and IFNγ treated neutrophils compared to WT, FDR<0.05, |logFC|>1). Interestingly, among the differentially expressed genes we identified several with established roles in AML biology. *HOXA9* and *HOXA10* were upregulated in NPs and monocytes, with *HOXA9* also upregulated in mature neutrophils. Overexpression of *HOXA9* and *HOXA10* has been linked to AML induction, maintenance, and adverse prognosis ^29,63,64^. Similarly, *FGF13*, whose expression is reduced in AML and is linked to poor overall survival ^65^, was found to be downregulated in *DNMT3A*-R882 mutant neutrophils and monocytes. Our data indicate that these transcriptional alterations are already present in CHIP, preceding malignant transformation. Gene Set Enrichment Analysis (GSEA) indicated that in the context of IFNγ stimulation, transcriptional signatures linked to inflammation, including those linked to IFN and TNF responses, were negatively enriched in *DNMT3A*-R882 monocytes compared to their WT counterparts. Nonetheless, *IL1A,* a key pro-inflammatory cytokine responsible for HSC functional attrition and inflammatory manifestations in a range of CHIP co-morbidities ^66,67^, was expressed at significantly higher levels in IFNγ-treated *DNMT3A*-R882 monocytes than WT monocytes. We conclude that *DNMT3A*-R882 mutations intrinsically induce alteration of monocyte inflammatory properties.

*DNMT3A*-R882 mutation also induced significant changes in mRNA expression along the neutrophil differentiation trajectory. GSEA analysis demonstrated that in untreated conditions, signatures linked to cell proliferation were enriched in *DNMT3A*-R882 NPs (**Fig.4D**) compared to WT NPs, in line with the increased neutrophil differentiation phenotype identified functionally for *DNMT3A*-R882 HSC/MPPs. In contrast, inflammatory signatures were significantly downregulated in *DNMT3A*-R882 NPs compared to their WT counterparts (**Fig.4D**). This shows that the attenuation of inflammatory signatures reported in *DNMT3A* mutant HSCs^42^ carries over during myeloid differentiation. However, *DNMT3A*-R882 mature neutrophils displayed increased maturity at the transcriptional level (**Fig.S4F**) and switched to a more pro-inflammatory transcriptional state than their WT counterparts when exposed to IFNγ (**Fig.4D**). A switch from dampened inflammatory transcriptional signatures of NPs to pro-inflammatory mature neutrophils was also observed in the *DNMT3A*-R882-edited mice compared to control mice (**Fig.4E**) in the xenotransplantation model (**Fig.S3C**). From this we conclude that while *DNMT3A*-R882 neutrophils progenitors display a dampened inflammatory profile at steady-state, their mature neutrophil progeny switches to a pro-inflammatory phenotype upon exposure to an inflammatory stimulus.

### Association of *DNMT3A*-R882 neutrophil*s* transcriptional signature to CHIP-related comorbidities

Neutrophils are known to contribute to pathogenesis due to their pro-inflammatory function in many diseases at increased risk in CHIP individuals, including chronic coronary syndrome (CCS) ^68,69^. We therefore sought to establish if *DNMT3A*-R882-induced transcriptomic alterations in mature neutrophils resemble neutrophil phenotypes found in CHIP comorbidities. First, we reanalysed a publicly available RNA-seq dataset of neutrophils isolated from CCS patients^70^. GSVA showed that the transcriptional signature associated with *DNMT3A*-R882 neutrophils (**Table S3**) was significantly enriched in neutrophils of CCS patients compared to controls (**Fig.5A**). Moreover, we found *DEFA3* to be among the most strongly upregulated genes in *DNMT3A*-R882 neutrophils (**Fig.5B** and **Fig.4D**). DEFA3 encodes a member of the neutrophil alpha-defensins (human neutrophil peptide, HNP) family, found abundantly in atherosclerotic plaques and proven to be highly atherogenic by promoting LDL transmural accumulation, monocyte endothelial recruitment, and macrophage to foam cell formation^71–73^.

**Fig. 5:**
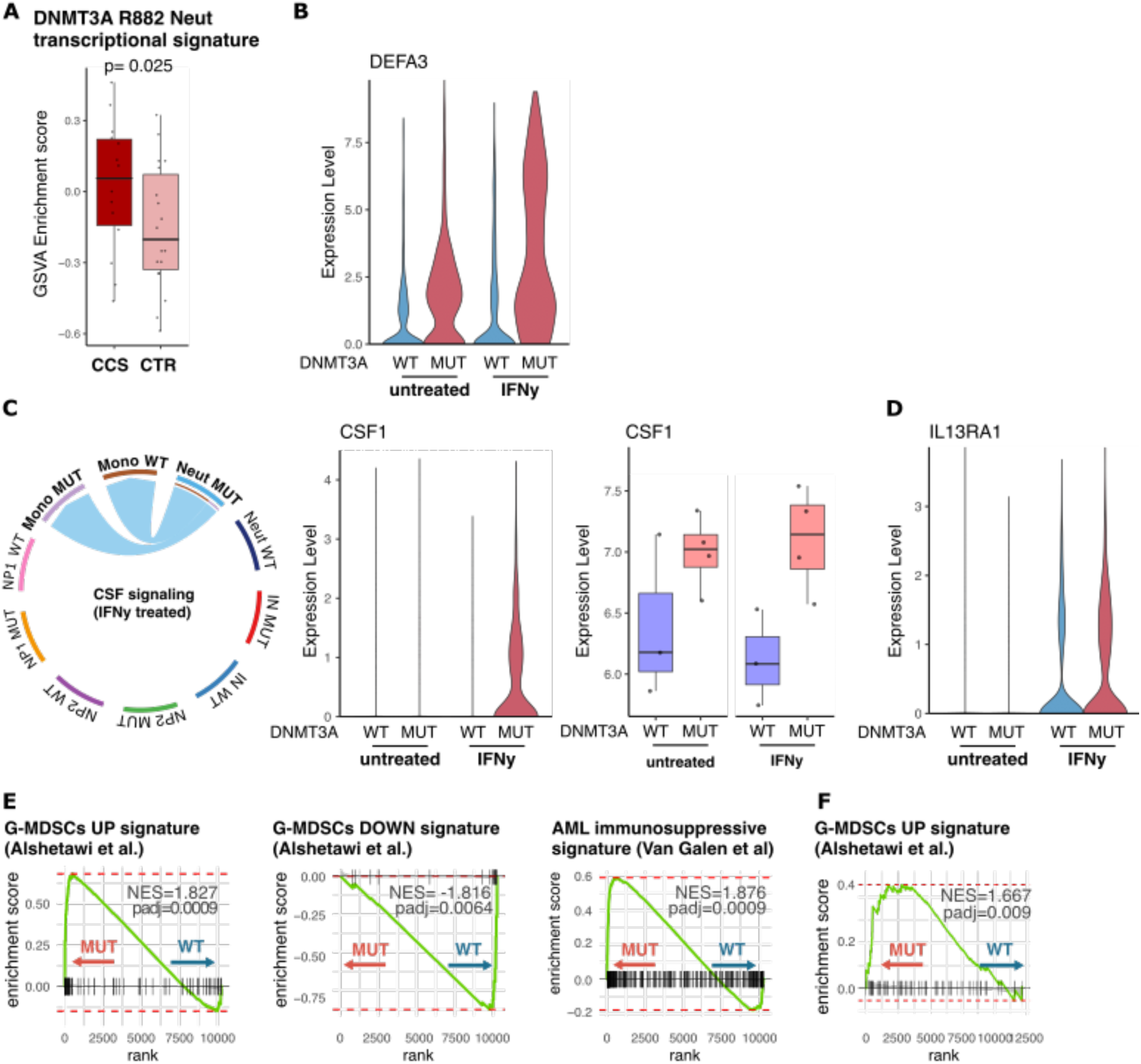
Transcriptomic signatures link *DNMT3A*-R882 mutant neutrophils to CHIP comorbidities. **(A)** GSVA enrichment score of the transcriptomic signature derived from *DNMT3A*-R882 CHIP neutrophils (from scRNAseq dataset, Fig4). Scores were calculated in bulk RNA-seq data of neutrophils from patients with chronic coronary syndrome (CCS, n=14) and control individuals (CTR, n=11)^70^. P value from linear modelling testing the association of disease group with GSVA score, adjusted for age. **(B)** Log-normalised expression levels of *DEFA3* gene in mature neutrophils from scRNA-seq. **(C)** Chord diagram of CSF soluble factor communication network inferred from ligand-receptor expression in scRNA-seq data using CellChat algorithm^74^ (left). Log-normalised expression levels of *CSF1* gene in mature neutrophils scRNA-seq (middle), and *CSF1* expression (log-TPM) in bulk RNA-seq of CRISPR-edited *in-vivo* HSC-derived neutrophils (NEU) and neutrophil precursors (PRE) (right). **(D)** Log-normalised expression levels of *IL13RA1* gene in scRNA-seq of mature neutrophils. **(E)** GSEA of immunomodulatory signatures in scRNA-seq of mature neutrophils under IFNγ stimulation. Granulocytic-MDSC (G-MDSC) signature from ^82^ and AML-linked myeloid immunosuppression signature from ^83^. **(F)** GSEA enrichment of G-MDSC immunomodulatory signature ^82^ in bulk RNA-seq of CRISPR-edited *in-vivo* HSC-derived neutrophils.

Following this finding, we further investigated whether mutant neutrophils establish aberrant, and potentially pathogenic, interactions with immune cell types. First, we interrogated our dataset employing the CellChat algorithm ^74^ to infer cell-to-cell interactions based on the expression of soluble mediators and receptors. This highlighted that under IFNγ stimulation, *DNMT3A*-R882 mutant neutrophils, but not WT neutrophils, signal to mutant and WT monocytes via the CSF network. Consistently, we found *CSF1* expression to be selectively upregulated in *DNMT3A*-R882 mutant neutrophils in scRNA-seq and in neutrophils derived *in vivo* from CRISPR *DNMT3A*-R882 edited HSCs (**Fig.5C**). CSF1 plays a central role in solid and haematological cancers, including AML, by inducing monocyte differentiation towards pro-tumorigenic M2 macrophages, hence promoting immune escape, cancer progression, and metastasis ^75–77^. Additionally, compared to WT neutrophils, *DNMT3A*-R882 neutrophils also overexpressed IL-13 receptor (**Fig.5D**), a central player in immunomodulatory pathways in which natural killer cells suppress tumour surveillance^78^. These observations indicate two potential mechanisms by which *DNMT3A*-R882 neutrophils may support tumour growth. Given the increased risk and adverse prognosis of CHIP in certain solid cancers, the finding that CH myeloid cells infiltrate the tumour microenvironment, and the growing interest in whether mutant leukocytes contribute to the pathogenesis of post-CHIP myeloid malignancy ^11,12,79,80^, we tested if *DNMT3A*-R882 neutrophils display transcriptional characteristics of myeloid derived suppressor cells (MDSCs). MDSCs are pathologically activated monocytes or neutrophils with immunosuppressive capacity, shown to contribute to cancer pathogenesis^81^ and poor outcomes in sepsis and an array of infectious diseases^62^. Interestingly, published signatures of MDSCs from breast cancer^82^ or AML^83^ patients were significantly enriched in IFNγ-exposed *DNMT3A-*R882 neutrophils (**Fig.5E**) and untreated *DNMT3A-*R882-edited neutrophils (**Fig.5F**) compared to WT. Collectively, these data indicate that *DNMT3A-*R882 neutrophils establish aberrant and most likely pathological interactions with other immune cells.

Overall, our findings show that while *DNMT3A-*R882 mutations impart a protective effect from inflammation during early stages of myeloid differentiation, they intrinsically determine the production of pro-inflammatory and immunosuppressive neutrophils akin to those produced during maladaptive responses ^62,92^ and found in non-haematopoietic CHIP comorbidities.

## Discussion

The impact of common CHIP mutations extends beyond the clonal advantage they impart to HSCs. In this study, we show that the *DNMT3A-*R882 mutation in human HSCs intrinsically alters myelopoiesis, in particular neutropoiesis, leading to remodelling of the innate immune repertoire. Mutant neutrophil progenitors differentiate rapidly and abundantly towards mature neutrophils, with potentiated pro-inflammatory and immunomodulatory transcriptional responses upon inflammatory challenge, akin to those observed in maladaptive myelopoiesis. Collectively, our study extends our understanding of how *DNMT3A-*R882 mutation predisposes to leukaemia and other CHIP co-morbidities.

Several findings from this study contribute to explain the increased risk of AML for *DNMT3A*-R882 carriers. First, our phylogenetic analysis showed increased clonal diversification in the *DNMT3A*-R882 clade, with more expansions of secondary drivers in the *DNMT3A*-R882 clade than primary drivers in the WT clade. Therefore, *DNMT3A*-R882 not only promotes clonal expansion of individual HSCs ^30,31,36^, but alters selective pressures facilitating the expansion of secondary mutations, eventually leading to clonal sweeps. A recent genomic study demonstrated that clonal somatic sweeps can occur up to decades before AML diagnosis and are a strong predictor of future AML ^84^. Second, less efficient monocyte production, with increased *HOXA* gene expression and decreased expression of several tumor suppressor genes, may seed the beginnings of the strong differentiation block observed in overt AML.

Both *TET2* deletion and *ASXL1* mutations have been recently shown to compromise neutrophil maturation^59,85^. Here we demonstrate the propensity of *DNMT3A-*R882 mutant HSC/MPPs to promote neutrophil maturation, while producing less monocytes. This occurs despite similar proportions of HSC/MPPs committing to these fates, suggesting that *DNMT3A-*R882 alters the kinetics of monocytic and neutrophil differentiation downstream of HSC/MPPs. These results functionally extend the observations of decreased monocyte precursors in *Dnmt3a*-null mouse HSCs^40^, and of higher neutrophil counts in individuals with germline^86,87^ or somatic^88^ *DNMT3A* mutations.

HSCs with *DNMT3A* mutations have been shown to preferentially expand in pro-inflammatory environment ^15,44^, as the one associated with aging and CHIP^18^. One explanation of this phenomenon is that *DNMT3A* mutations confer an anti-inflammatory transcriptional state ^42,89^, which protects HSCs from the negative long-term impact of chronic inflammation ^14,17^. Here we demonstrate that *DNMT3A-* R882 mutations promote accelerated differentiation of neutrophil progenitors that, like *DNMT3A-* R882 HSCs, are intrinsically resilient to inflammation, compared to their WT counterparts in the same individual. However, mature *DNMT3A-*R882 neutrophils switch to a pro-inflammatory state, particularly upon exposure to inflammatory stimuli. This identifies *DNMT3A-*R882 neutrophils as contributors to the self-perpetuating inflammatory circle that favours the clonal expansion of mutated HSCs in CHIP.

Mature innate immune cells, such as neutrophils, monocytes, macrophages or NK cells, gain enhanced responses after their first encounter with a specific immune stimulus, in a phenomenon termed “trained immunity”. This memory is established and maintained at HSPC level ^90^. Trained immunity is usually protective, both in the context of infections and cancer, but becomes deleterious (“maladaptive”) in chronic inflammatory conditions or in specific subsets of patients ^62,91–93^. Here we demonstrate that *DNMT3A-*R882 mutations intrinsically induce dysfunctional innate immune cell differentiation trajectories in humans in a manner that i) exacerbates production of inflammatory cytokines such as IL1a (produced by monocytes) and IL1b and TNF (produced by neutrophils), ii) elicits immunosuppressive phenotypes in mature neutrophils; iii) induces broader immune remodelling likely extending immune dysregulation from neutrophils to macrophages, NK cells and T cells. While we cannot exclude that some of the functions mediated by *DNMT3A-*R882 neutrophils may be beneficial, several pieces of evidence point towards an overall maladaptive effect. In line with a study that involved *Dnmt3a-*R887H neutrophils into the bone loss associated with periodontitis and arthritis in mouse models^8^, we find that neutrophils differentiated from *DNMT3A-*R882 HSC/MPPs of CHIP individuals acquire transcriptomic signatures similar to those found in patients with chronic coronary syndromes. Moreover, *DNMT3A-*R882 neutrophils upregulated transcriptional features of myeloid-derived suppressor cells, which are known to promote immune evasion in solid tumours ^81^.

We propose that this intrinsic maladaptive immunity feature of *DNMT3A-R882* mutations is a key player in determining susceptibility of *DNMT3A* CHIP carriers to develop a wide-range of non-haematological CHIP co-morbidities, ranging from inflammatory diseases to solid tumours. This is of clinical relevance as models to predict the risk of progression towards malignancy or cardiovascular events for CHIP carriers are emerging, together with potential therapeutic strategies for early intervention targeting specific somatic clones.

### Limitations

Our study was conducted on primary samples carrying *DNMT3A-*R882 mutations at relatively high VAF, which are rare in the general population. To mitigate for the relatively small sample size (10 individuals), we have consistently compared WT and *DNMT3A-*R882 phenotypes within the same individual. We have also validated relevant phenotypes using xenotransplantation of HSC/MPPs where *DNMT3A-R882* was introduced by gene editing. Whereas this design is sufficiently powered to identify features of myeloid differentiation associated with *DNMT3A-*R882, it is not powered to identify potential differences between *DNMT3A-*R882H and R882C. We did not note any phenotypic difference between *DNMT3A-*R882H and R882C in this study. To date, these two variants have been reported to possess similar hypomethylation patterns^23^, functional effects in cell lines^94^ and impart similar clinical outcomes in AML patients^95^.

The myeloid cells analysed in this study were derived either *in vitro* from human HSC/MPPs isolated from *DNMT3A-*R882 carriers, or *in vivo* from HSC/MPPs in which *DNMT3A-*R882 was introduced via CRISPR editing. Prior isolation of mononuclear cells and/or the fragility of neutrophils through the freeze/thaw process prevented the analysis of neutrophils directly isolated from *DNMT3A-*R882 carriers. Nonetheless this differentiation approach offers the opportunity to directly compare the dynamics of differentiation between isogenic WT and *DNMT3A*-R882 HSC/MPPs in a controlled environment, therefore revealing properties that would be missed by snapshot analyses of differentiated cells.

## STAR Methods

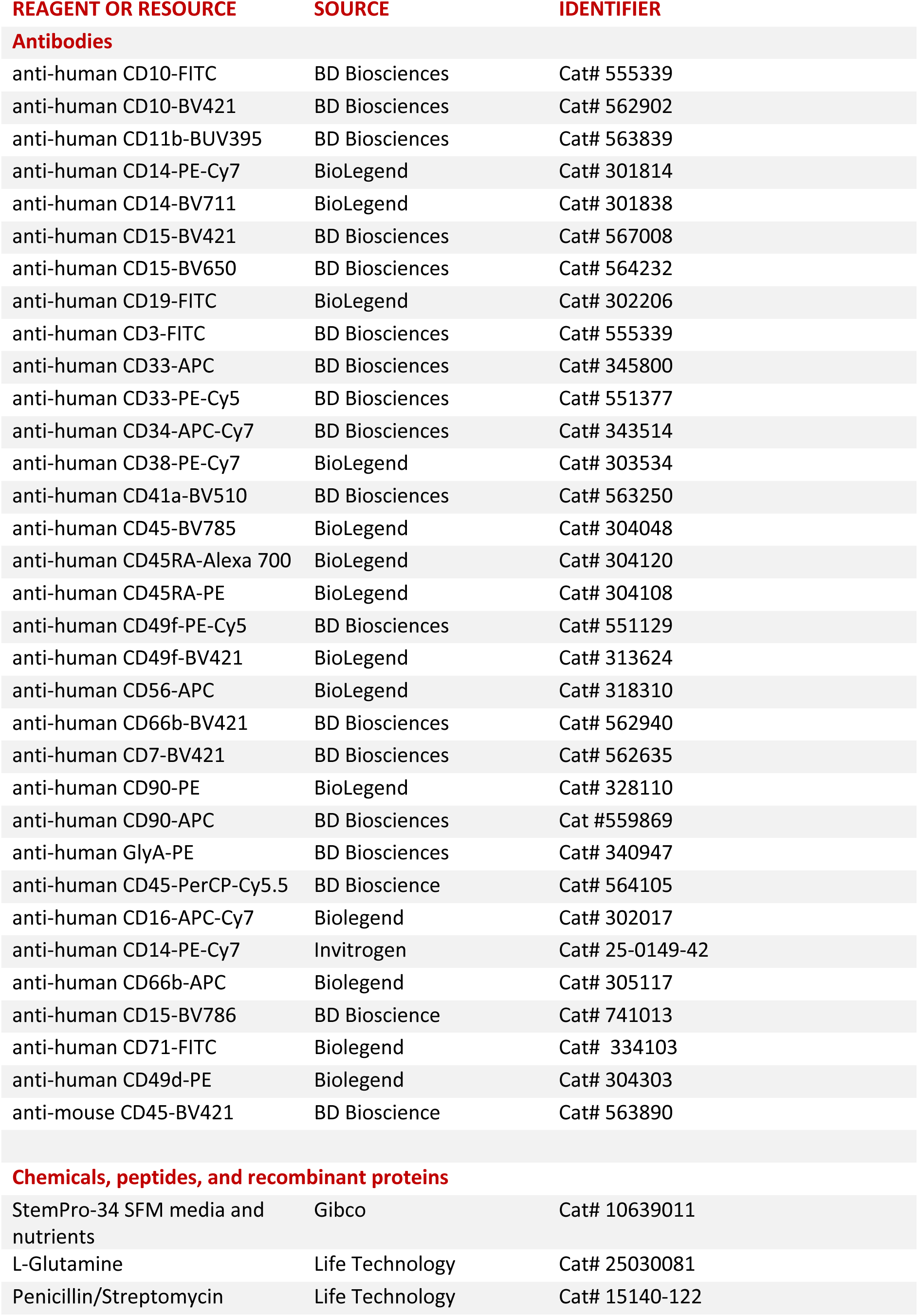
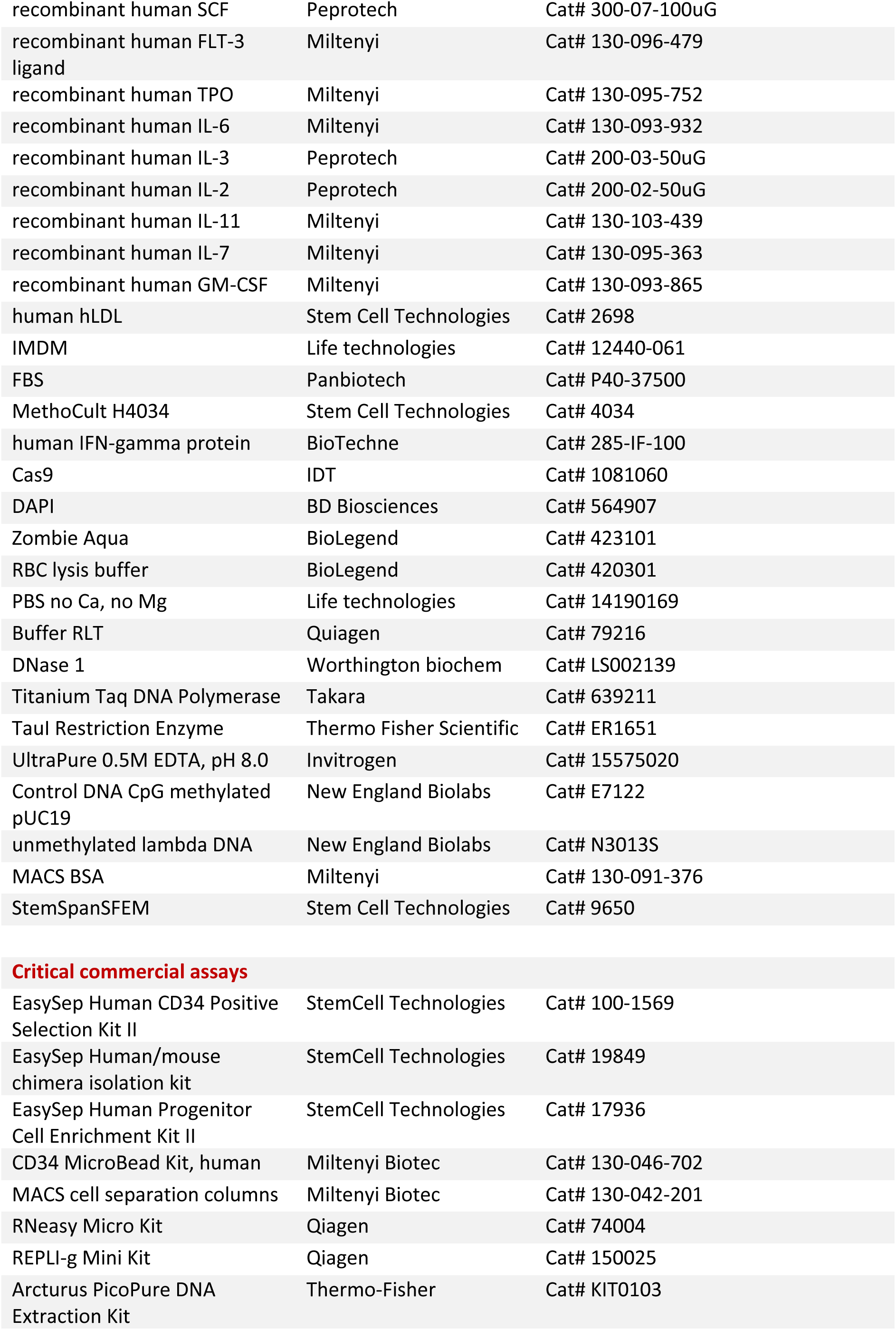
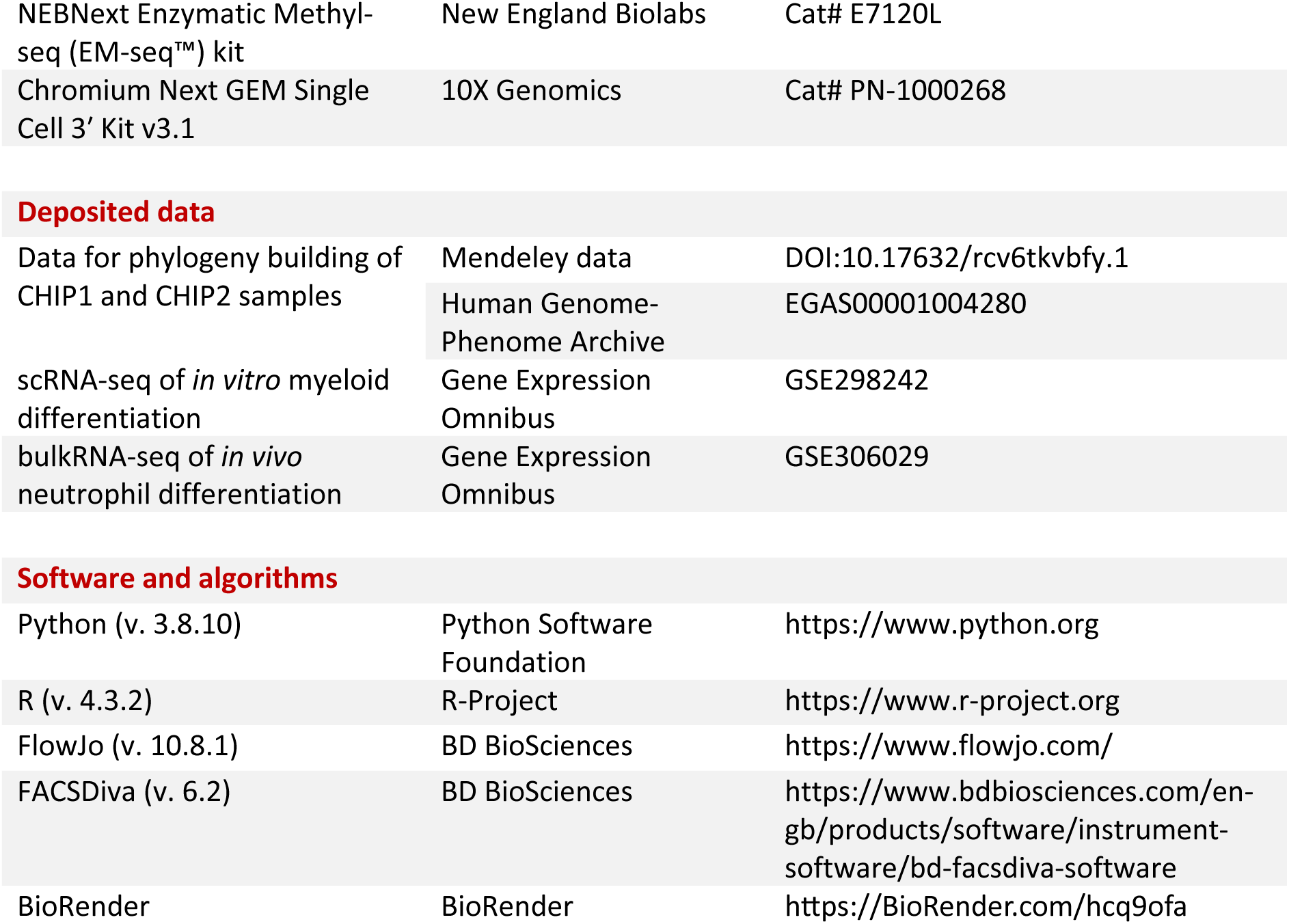
KEY RESOURCE TABLE.

KEY RESOURCE TABLE
| REAGENT OR RESOURCE | SOURCE | IDENTIFIER |
| --- | --- | --- |
| **Antibodies** |  |  |
| anti-human CD10-FITC | BD Biosciences | Cat# 555339 |
| anti-human CD10-BV421 | BD Biosciences | Cat# 562902 |
| anti-human CD11b-BUV395 | BD Biosciences | Cat# 563839 |
| anti-human CD14-PE-Cy7 | BioLegend | Cat# 301814 |
| anti-human CD14-BV711 | BioLegend | Cat# 301838 |
| anti-human CD15-BV421 | BD Biosciences | Cat# 567008 |
| anti-human CD15-BV650 | BD Biosciences | Cat# 564232 |
| anti-human CD19-FITC | BioLegend | Cat# 302206 |
| anti-human CD3-FITC | BD Biosciences | Cat# 555339 |
| anti-human CD33-APC | BD Biosciences | Cat# 345800 |
| anti-human CD33-PE-Cy5 | BD Biosciences | Cat# 551377 |
| anti-human CD34-APC-Cy7 | BD Biosciences | Cat# 343514 |
| anti-human CD38-PE-Cy7 | BioLegend | Cat# 303534 |
| anti-human CD41a-BV510 | BD Biosciences | Cat# 563250 |
| anti-human CD45-BV785 | BioLegend | Cat# 304048 |
| anti-human CD45RA-Alexa 700 | BioLegend | Cat# 304120 |
| anti-human CD45RA-PE | BioLegend | Cat# 304108 |
| anti-human CD49f-PE-Cy5 | BD Biosciences | Cat# 551129 |
| anti-human CD49f-BV421 | BioLegend | Cat# 313624 |
| anti-human CD56-APC | BioLegend | Cat# 318310 |
| anti-human CD66b-BV421 | BD Biosciences | Cat# 562940 |
| anti-human CD7-BV421 | BD Biosciences | Cat# 562635 |
| anti-human CD90-PE | BioLegend | Cat# 328110 |
| anti-human CD90-APC | BD Biosciences | Cat #559869 |
| anti-human GlyA-PE | BD Biosciences | Cat# 340947 |
| anti-human CD45-PerCP-Cy5.5 | BD Bioscience | Cat# 564105 |
| anti-human CD16-APC-Cy7 | Biolegend | Cat# 302017 |
| anti-human CD14-PE-Cy7 | Invitrogen | Cat# 25-0149-42 |
| anti-human CD66b-APC | Biolegend | Cat# 305117 |
| anti-human CD15-BV786 | BD Bioscience | Cat# 741013 |
| anti-human CD71-FITC | Biolegend | Cat# 334103 |
| anti-human CD49d-PE | Biolegend | Cat# 304303 |
| anti-mouse CD45-BV421 | BD Bioscience | Cat# 563890 |
| **Chemicals, peptides, and recombinant proteins** |  |  |
| StemPro-34 SFM media and nutrients | Gibco | Cat# 10639011 |
| L-Glutamine | Life Technology | Cat# 25030081 |
| Penicillin/Streptomycin | Life Technology | Cat# 15140-122 |
| recombinant human SCF | Peprotech | Cat# 300-07-100uG |
| recombinant human FLT-3 ligand | Miltenyi | Cat# 130-096-479 |
| recombinant human TPO | Miltenyi | Cat# 130-095-752 |
| recombinant human IL-6 | Miltenyi | Cat# 130-093-932 |
| recombinant human IL-3 | Peprotech | Cat# 200-03-50uG |
| recombinant human IL-2 | Peprotech | Cat# 200-02-50uG |
| recombinant human IL-11 | Miltenyi | Cat# 130-103-439 |
| recombinant human IL-7 | Miltenyi | Cat# 130-095-363 |
| recombinant human GM-CSF | Miltenyi | Cat# 130-093-865 |
| human hLDL | Stem Cell Technologies | Cat# 2698 |
| IMDM | Life technologies | Cat# 12440-061 |
| FBS | Panbiotech | Cat# P40-37500 |
| MethoCult H4034 | Stem Cell Technologies | Cat# 4034 |
| human IFN-gamma protein | BioTechne | Cat# 285-IF-100 |
| Cas9 | IDT | Cat# 1081060 |
| DAPI | BD Biosciences | Cat# 564907 |
| Zombie Aqua | BioLegend | Cat# 423101 |
| RBC lysis buffer | BioLegend | Cat# 420301 |
| PBS no Ca, no Mg | Life technologies | Cat# 14190169 |
| Buffer RLT | Quiagen | Cat# 79216 |
| DNase 1 | Worthington biochem | Cat# LS002139 |
| Titanium Taq DNA Polymerase | Takara | Cat# 639211 |
| TauI Restriction Enzyme | Thermo Fisher Scientific | Cat# ER1651 |
| UltraPure 0.5M EDTA, pH 8.0 | Invitrogen | Cat# 15575020 |
| Control DNA CpG methylated pUC19 | New England Biolabs | Cat# E7122 |
| unmethylated lambda DNA | New England Biolabs | Cat# N3013S |
| MACS BSA | Miltenyi | Cat# 130-091-376 |
| StemSpanSFEM | Stem Cell Technologies | Cat# 9650 |
| Critical commercial assays |  |  |
| EasySep Human CD34 Positive Selection Kit II | StemCell Technologies | Cat# 100-1569 |
| EasySep Human/mouse chimera isolation kit | StemCell Technologies | Cat# 19849 |
| EasySep Human Progenitor Cell Enrichment Kit II | StemCell Technologies | Cat# 17936 |
| CD34 MicroBead Kit, human | Miltenyi Biotec | Cat# 130-046-702 |
| MACS cell separation columns | Miltenyi Biotec | Cat# 130-042-201 |
| RNeasy Micro Kit | Qiagen | Cat# 74004 |
| REPLI-g Mini Kit | Qiagen | Cat# 150025 |
| Arcturus PicoPure DNA Extraction Kit | Thermo-Fisher | Cat# KIT0103 |
| NEBNext Enzymatic Methyl-seq (EM-seq™) kit | New England Biolabs | Cat# E7120L |
| Chromium Next GEM Single Cell 3' Kit v3.1 | 10X Genomics | Cat# PN-1000268 |
| Deposited data |  |  |
| Data for phylogeny building of CHIP1 and CHIP2 samples | Mendeley data | DOI:10.17632/rcv6tkvbfy.1 |
|  | Human Genome-Phenome Archive | EGAS00001004280 |
| scRNA-seq of *in vitro* myeloid differentiation | Gene Expression Omnibus | GSE298242 |
| bulkRNA-seq of *in vivo* neutrophil differentiation | Gene Expression Omnibus | GSE306029 |
| Software and algorithms |  |  |
| Python (v. 3.8.10) | Python Software Foundation | https://www.python.org |
| R (v. 4.3.2) | R-Project | https://www.r-project.org |
| FlowJo (v. 10.8.1) | BD BioSciences | https://www.flowjo.com/ |
| FACSDiva (v. 6.2) | BD BioSciences | https://www.bdbiosciences.com/en-gb/products/software/instrument-software/bd-facsdiva-software |
| BioRender | BioRender | https://BioRender.com/hcq9ofa |

## RESOURCE AVAILABILITY

**Lead contact:** any requests for resources and/or reagents should be directed to the corresponding author, Elisa Laurenti.

**Materials availability:** no new reagents were generated in this study.

### Data and code availability

RNA sequencing data have been deposited at GEO under accession numbers GSE298242 (https://www.ncbi.nlm.nih.gov/geo/query/acc.cgi?acc=GSE298242) and GSE306029 (https://www.ncbi.nlm.nih.gov/geo/query/acc.cgi?acc=GSE306029). The data needed to reproduce the phylogenetic analysis of CHIP1 and CHIP2 are publicly available at European Genome-phenome Archive (https://ega-archive.org/studies/EGAS00001004280) and Mendeley Data at https://data.mendeley.com/datasets/rcv6tkvbfy/1.

All code is available at https://github.com/laurenti-lab/DNMT3A_CH.

## EXPERIMENTAL MODELS AND STUDY PARTICIPANT DETAILS

### Patient sample selection and ethics approval

Primary patient PB/mPB samples were received from Princess Margaret Cancer Centre, University Health Network (UHN), Canada (UHN IRB protocol 01-0573) and from Rambam Health Care Campus, Israel (IRB protocol #283-1) via the Weizmann Institute for Medical Science, Newcastle Hospitals NHS Foundation Trust (Newcastle Biobank REC reference 17/NE/0361), from the SardiNIA longitudinal cohort study (Sardinian Regional Ethics Committee prot. n. 2171/CE), and from the Cambridge Blood and Stem Cell Biobank study (REC ID 24/EE/0116, previously 18/EE/0199). All PB/mPB/BM samples selected for the study had a *DNMT3A*-R882 VAF >2% and were collected after informed consent.

Human HSCs were isolated from umbilical cord blood (UCB) samples collected from full term donors after informed consent and obtained from the Anthony Nolan charity.

Frozen MNCs were originally sourced from G-CSF-mobilised (mPB) and non-mobilised peripheral blood (PB) or BM as specified in **Table S1**. PB samples were collected by phlebotomy, mPB by apheresis and BM by aspiration. MNCs were isolated following Ficoll-Paque separation (GE 67 Healtcare, London) and viably frozen with or without RBC lysis (RBC lysis buffer, Biolegend).

All work in Cambridge was performed under the Cambridge Blood and Stem Cell Biobank REC approval 18/EE/0199 and 24/EE/0116. All work in London was performed under Crick HTA licence 12650.

In this work we performed two sets of animal experiments, with two different mouse strains. For all experiments with “NBSGW-SGM3” mice, these were produced at the Francis Crick Institute by breeding for 7 generations NBSGW and NSG-SGM3 (NOD. Prkdcscid Il2rgtm1 Tg(CMV-IL3, CSF2, KITLG). Mice were bred in isolators with aseptic standard operating procedures in the Biological Research Facility of The Francis Crick Institute. Once weaned, mice were kept in ventilated cages. All animal experiments were performed under the U.K Home Office project license (70/8904) in accordance with The Francis Crick Institute animal ethics committee guidance.

For all AML xenotransplantation experiments, assays were performed at the Weizmann Institute, in accordance with the institutional guidelines approved by the Weizmann Institute of Science Animal Care Committee (11790319-2).

## METHOD DETAILS

### HSC/MPP isolation

#### CD34+ enrichment

Samples from CHIP donors and AML patients were received as frozen MNCs. MNCs from each individual were thawed using 50% IMDM/50% FCS with 1:100 DNase. For samples with >5×10^7^ MNCs, positive selection of CD34+ cells from the remaining sample was achieved by incubating with CD34 Micro Beads (30µL/10^8^ cells, Miltenyi), FcR Blocking Reagent (30µL/10^8^ cells) and PBS + 3% FCS (90µL/10^8^ cells) for 30 minutes at 4°C. Cells were then washed and resuspended in MACS buffer and applied to a prepared LS magnetic column for manual selection as per the Miltenyi user manual.

Samples with 1-5×10^7^ MNCs were selected using EasySep Human CD34 Positive Selection Kit II (StemCell Technologies), according to manufacturer instructions. Isolated CD34+ cells underwent staining and fluorescence-activated cell sorting (FACS). Samples with < 1 x 10^7^ MNCs after thawing were processed for FACS without prior magnetic selection.

For CB samples, lineage marker positive cells were depleted using an EasySep Human Progenitor Cell Enrichment Kit (Stem Cell Technologies).

#### Flow cytometry sorting of HSC/MPPs

Single HSC/MPPs from CHIP donors and AML patients were sorted directly into 96 well plates containing 100µL/well MEM media: StemPro base media supplemented with Nutrients, Pen/Strep (1%), L-Glu (1%), human LDL (50ng/ml), rhSCF (100ng/ml), rhFlt-3L (20ng/ml), rhTPO (100ng/ml), EPO (3 units/ml), rhIL-6 (50ng/ml), rhIL-3 (10ng/ml), rhGM-CSF (20ng/ml), rhIL-11 (50ng/ml), rhIL-2 (10ng/ml), and rhIL-7 (20ng/ml).

Sorting gates used were as follows: Cells/singlets/live cells/CD19-/CD33-/CD34+/CD45dim/CD38-/CD45RA-or Lin-(CD3/CD19/CD20/CD14/CD56/CD11c)/CD33-/CD34+/CD45dim/CD38-/CD45RA-. For CHIP1, GMPs were also sorted using the above strategy until CD45dim, followed by CD38+/CD10-/CD7-/CD45RA+ (**Fig. S1A**).

Antibodies staining was performed in 100µL for 20min at room temperature (RT) with the following panels:

*CHIP1*: CD33 APC/CD45RA AF-700/CD34 APC-Cy7/CLEC9A PE/CD49f PE-Cy5/CD38 PE-Cy7/CD90 FITC/CD10 BV421/CD7 BV421/Zombie Aqua BV510/CD45 BV605/CD19 BV785

*CHIP3, CHIP4, AML1, AML2, AML3, AML4, AML5*: CD33 APC/CD45RA AF-700/CD34 APC-Cy7/CD90 PE/CD49f PE-Cy5/CD38 PE-Cy7/CD19 FITC/CD10 BV421/CD7 BV421/Zombie Aqua BV510/ CD45 BV785

*CHIP5*: CD49f BV421/ Zombie Aqua BV510/ CD19 FITC/ CD45RA PE/ CD38 PE-Cy7/ CD90 APC/ CD34 APC-Cy7

*CHIP6*: CD33 APC/CD45RA AF-700/CD34 APC-Cy7/CD90 PE/CD49f PE-Cy5/CD38 PE-Cy7/Lineage (CD19, CD3, CD14, CD56, CD20) FITC/CD11c FITC/CD10 BV421/CD7 BV421/Zombie Aqua BV510/CD45 BV785

HSC/MPPs or GMPs were sorted on FACS Aria Fusion (BD) or Influx (BD).

Sorting of Lin-/CD34+/CD38-for CRISPR editing was performed as previously described ^96^.

### Single cell culture and phenotyping

Single HSC/MPPs or GMPs were cultured using sterile technique and all wells containing visible colonies were harvested after 3 weeks in culture. Media was added to 20-50% of wells in each experiment as needed, on average once over the 21-day culture period. At the time of analysis, all colonies were harvested into a new 96 well plates and split 2/3 for flow cytometry phenotyping analysis and 1/3 for genotyping. Colonies for phenotyping were stained with using 50µL/well antibody panel for 20 minutes at RT. Antibodies used are listed in the key resource table. Panels used were as follows:

*CHIP1*: CD56 APC/CD11b APC-Cy7/GlyA PE/CD45 PE-Cy5/CD14 PE-Cy7/CD41 FITC/CD15 BV421/CD41 FITC

*CHIP4, AML2, AML3:* CD3 APC-Cy7/CD19 APC-Cy7/CD10 APC-Cy7/CD71 PE/CD45 PE-Cy5/GlyA PE-Cy7/CD56 FITC/CD66b BV421/CD15 BV650/CD14 BV711/CD11b BUV395

*AML1:* CD56 APC/CD14 AF-700/CD34 APC-Cy7/GlyA PE/CD33 PE-Cy5/CD38 PE-Cy7/CD16 FITC/CD15 BV421/CD45 BV510

*AML4:* CD56 APC/GlyA PE/CD3 FITC/CD19 FITC/CD10 FITC/CD66b BV421/CD15 BV510/CD45 BV605/CD14 BV711/CD11b BUV395

After washing, colonies were resuspended in 100µL PBS + 3% FCS and analysed using a LSR Fortessa (BD) on HTS mode (50 μl extraction. All remaining cells from each colony were frozen in 50µL PBS f for future DNA extraction.

For CHIP2, Peripheral blood mononuclear cells were isolated from blood and plated at 50,000 cells per ml in MethoCult H4034 (Stemcell Technologies). After 14 days in culture, 96 single haematopoietic colonies were plucked per individual (total 288 colonies, each made up of hundreds to thousands of cells) and lysed in 50 μl of RLT lysis buffer (Qiagen).

### DNA extraction and sequencing

#### Initial targeted screening for selection of primary samples

CHIP and AML patients’ samples were sequenced at the Weizmann Institute of Science, Israel, as published in Tuval et al. ^97^. DNA from CD3 depleted (or CD34+ enriched) cells from primary PB samples was extracted using the Qiagen DNeasy Blood and Tissue kit as per the manufacturer’s protocol. Samples with low viable cell numbers had DNA extraction by a different method: centrifugation of 50,000 cells, lysis by incubation with 50µL NaOH 50mM at 99°C for 10min, cooling to RT and addition of 5µL Trs 1M pH8; 4.5µL of this solution was taken for library preparation. Libraries were prepared for Next Generation Sequencing (NGS) using single molecule Molecular Inversion Probes (smMIPs) ^98,99^, designed with MIPgen software. The final smMIP panel was designed to identify recurrently mutated AML hotspots in 33 genes as previously outlined in Tuval et al ^97^. DNA was sequenced in duplicates directly following extraction without whole genome amplification.

#### Targeted sequencing of single cell derived colonies

For samples CHIP1, CHIP3, CHIP4, CHIP6, AML1, AML2, AML3 and AML4, DNA was extracted and amplified using the Qiagen RepliG amplification kit as per manufacturer’s protocol. An amplicon-based approach was used to generate libraries to sequence for targeted mutations including DNMT3A R882, NPM1 and FLT3-ITD. Following 2 PCR amplification cycles, samples were pooled. PCR purification was then performed using DNA clean and concentrator-5, and library evaluation was done on TapeStation as per manufacturer’s protocol. Cleaning was performed by Blue Pippin and Qubit 4 Fluorometer was used to measure library concentration of each well using iQuant dsDNA Assay kit as per manufacturer’s protocol.

Primary PCR for DNMT3A R882 (exon 23) was performed with the following primers:

Forward: CTACACGACGCTCTTCCGATCTTAACTTTGTGTCGCTACCTC

Reverse: CAGACGTGTGCTCTTCCGATCTTTTTCTCCCCCAGGGTATTTG

Secondary PCR was performed with the following primers:

Forward: AATGATACGGCGACCACCGAGATCTACAC [Fw_Index_D5XX] ACACTCTTTCCCTACACGACGCTCTTCCG;

Reverse primer:

CAAGCAGAAGACGGCATACGAGAT [Rev_Index_D7XX]GTGACTGGAGTTCAGACGTGTGCTCTTCCG

All sequencing steps performed at the Weizmann institute used either MiSeq, MiniSeq or NovaSeq sequencers (Illumina).

#### Rapid genotyping of single cell derived colonies

For CHIP5, half of every single cell derived colony was harvested at day 17 for DNA extraction, while the rest was left in culture for a total of three weeks. DNA was extracted using the Arcturus PicoPure kit (Thermo Fisher Scientific): cells were pelleted and resuspended in 17μl of Proteinase K buffer, prior of incubation in a G-Storm thermocycler (65°C for 6h, 75°C for 30min).

As the presence of the DNMT3A c.2645 p.R882 mutation ablates the binding site of TauI restriction enzyme ^100^, we employed a restriction assay to genotype the colonies. The region of interest was amplified with TitaniumTaq Polymerase (Takara Bio) in a two-step PCR reaction for 33 cycles (30 seconds at 95°C, 3minutes at 68 °C, primers F: GTGTGGTTAGACGGCTTCCG, R: ACAAGAGGTAACAGCGGCTT) generating a PCR product of 997bp, which encompasses the c.2645 mutation site and an additional TauI binding site, serving as internal control of the reaction. 10 μl of PCR product were digested with 0.9 μl of TauI (3U/μl, Thermo Fisher Scientific) and 2.3 μl of restriction buffer B (Thermo Fisher Scientific). The digestion was performed at 55 °C for an hour and stopped with 0.8 μl of EDTA (0.5M, Thermo Fisher Scientific). Reaction products were run on a 1.5% agarose gel: WT PCR products cut into 691bp fragments, DNMT3A c.2645 mutant ones into 813 bp fragments, while uncut products remained 997bp long. Genotyping was confirmed with Sanger sequencing of PCR products from randomly picked colonies (Sequencing primer S: CACACACGCAAAATACTCC), showing 100% concordance between the genotyping and sequencing results.

#### Whole genome sequencing (WGS)

DNA from 127 single HSPC derived colonies from CHIP1 was extracted using the Arcturus PicoPure DNA extraction kit (Thermo Fisher Scientific). Colonies were transferred to a PCR plate, pelleted, and resuspended in 17µL Proteinase K buffer per well. Plates were incubated on a G-Storm thermocycler at 65°C for 6h, 75°C for 30min. For CHIP2, 96 single MNC derived single cell colonies were lysed in 50 ul of RLT lysis buffer (Qiagen). Library preparation for WGS was performed using the low-input pipeline at the Wellcome Trust Sanger Institute as previously described ^101^. 150 bp paired-end sequencing reads were generated for CHIP1 on the Hiseq X platform (Illumina) at an average sequencing depth of 13X and for CHIP2 on the NovaSeq 6000 platform (Illumina) with average sequencing depth of 15X.

#### Whole genome methylation sequencing (EM-seq)

Whole genome methylation sequencing library preparation was performed using the NEBNext Enzymatic Methyl-seq (EM-seq™) kit (New England Biolabs) on DNA from colonies of CHIP1. To assess conversion efficacy, CpG-methylated pUC19 (New England Biolabs) was included as a positive control and unmethylated lambda DNA (New England Biolabs) as a negative control, both added prior to enzymatic conversion and library preparation. 150 bp paired-end sequencing reads were generated on the NovaSeq 6000 platform (Illumina), with average sequencing depth of 15.5X.

#### scRNA-seq and IFNγ in vitro challenge

Single-cell transcriptomic analysis was performed on single HSC/MPP derived colonies from one CHIP individual, CHIP5. After 17 days of culture, half of each colony was harvested for genotyping. After three weeks of culture, randomly selected single cell derived colonies were split in two and one half of every colony underwent overnight stimulation with IFNγ (50 ng/ml of recombinant human IFN-gamma protein, Bio-Techne). A total of 30 colonies were harvested, stained with fluorescence conjugated antibodies (CD45 FITC, CD11b APC-Cy7, CD14 PE-Cy7, GlyA PE, CD66b APC, BV421 CD15, Zombie Aqua) at RT for 20 minutes. Live myeloid cells (Zombie-/GlyA-), neutrophils (Zombie-/GlyA-/CD14-/CD66b+/CD15+), and monocytes (Zombie-/GlyA-/CD14+/CD11b+) were sorted on Influx cell sorter (BD) with a 140 μm nozzle (**Fig.S2B**). Single cell sequencing libraries were generated on the Chromium 10X platform (10X Genomics) using the v3.1 kit according to manufacturer’s instructions. Time from sort to encapsulation was less than 1 hour. Cells suspensions (in PBS containing 0.04% BSA (400μg/ml)) were kept at 4°C while awaiting to be processed.

#### Introduction of DNMT3A-R882H mutation in primary human HSCs and generation of humanized mice reconstituted with CRISPR-edited hHSCs

After sorting (see above), HSCs (Lin-/CD34+/CD38-) were seeded in StemSpanSFEM (Stem Cell Technologies) supplemented with 100 ng/mL rhFLT-3L, 100 ng/mL rhSCF and 100 ng/mL rhTPO (Thermofisher-Peprotech, Cat# 300-19-10UG, 300-07-10UG and 300-18-10UG respectively). Introduction of R882H mutation in the *DNMT3A* gene by CRISPR was done following our previous protocol ^37^. Human HSCs were collected for CRISPR editing after 48 hours from the cell sorting. For CRISPR editing we used the NEON Transfection system (Thermofisher). After the electroporation cells were cultured in the same media for 48 hours before being injected into “NBSGW-SGM3” mice (10,000-20,000 Lin-CD34+CD38-cells/mouse) via intravenous administration. Engraftment and mutation efficiency within the human hematopoietic system reconstituted were analysed for each mouse by bone marrow aspiration at 6– and 9-weeks post injection, and the mice were sacrificed between 12-14 weeks post-transplantation. *DNMT3A*-R882H VAF at the final time point was analysed for each mouse by next generation sequencing (results in **Table S4**). All humanized mice included in this study had >60% human haematopoietic cell engraftment in the bone marrow.

#### Xenotransplantation for selection of AML samples to be used in this study

Primary CD3 depleted PB samples from patients with AML were injected into SGM3, NSG or hSCF mice to assess engraftment capacity as detailed in ^97^. Engrafted human cells were then assessed by flow cytometry; engraftment was defined multi-lineage if ≥10% cells were non-myeloid/lymphoid (CD19+) and both myeloid and lymphoid cells were observed. This confirms non-leukemic engraftment as per Shlush et al ^36^. All AML samples used in this study engrafted multilineage.

## QUANTIFICATION AND STATISTICAL ANALYSIS

### Data processing and variant calling for DNMT3A targeted sequencing

For samples CHIP3, CHIP4, CHIP6, AML1, AML2, AML3, AML4 and AML5, paired-end 2X151bp sequencing data were converted to fastq format. Reads were merged using BBmerge v38.62 ^102^ with default parameters, followed by trimming of the ligation and extension arm using Cutadapt v2.10 ^103^. Unique Molecular Identifiers were trimmed and assigned to each read header. Processed reads were aligned using BWA-MEM ^104^ to a custom reference genome, comprised of the appropriate smMIP panel sequences ±150 bases extracted from GRCh37 hg19. Aligned files were sorted, converted to BAM followed by Indel realignment using IndelRealigner (GATK v.3.7 ^105^). Genotype analysis was based on targeted sequencing data using 3 independent variant callers: MuTect2 in a tumor-only mode, Varscan2 v2.3.9 ^106^ and Platypus ^107^ for indels.

Colonies with adequate sequencing depth were assessed using all 3 variant callers to determine VAF and classified as WT (<2% VAF) or *DNMT3A*-R882 (≥2% VAF). Prior to this step, colonies with low sequencing depth were filtered, to avoid the mis-assignment of low-sequencing depth results to WT calls. Per run, the minimum sequencing depth threshold was selected as the minimum depth associated with the detection of ≥2% VAF in at least two variant callers. Colonies below this threshold were excluded. Variant calling results for this study are shown in **Table S2.** Genotyping data was then correlated with the phenotyping data available from flow cytometry.

To assess the level of CRISPR editing, human CD45+ cells were purified from xenografted mice either by flow sorting or by isolating hCD45+ with magnetic beads (EasySep Human/mouse chimera isolation kit, Stem Cell Technologies). Cells were then centrifuged and kept at –20C until DNA extraction. PCR was performed to amplify the *DNMT3A* region using the following primers:

fwd: TCGTCGGCAGCGTCAGATGTGTATAAGAGACAGAGGAGTTGGTGGGTGTGAGT, rev: GTCTCGTGGGCTCGGAGATGTGTATAAGAGACAGCACGCAAAATACTCCTTCAGC.

The PCR product was then sequenced with MiSeq.

### CytoTree

In an *in silico* pooled analysis approach, we pooled all mature from all colonies within each individual sample without any downsampling. Data from the live “single cells” gate of each colony (**Fig.S1B**) within each individual sample were exported as .fcs files. Using CytoTree, a pre-existing R/Bioconductor package ^56^, .fcs files from all colonies derived from each donor were pooled and converted into a cell expression matrix for selected markers of mature blood cell production (CD45, GlyA, CD56, CD11b, CD15, CD14 and CD66b where available). No normalization was performed at this step. Metadata such as cell genotype (WT or *DNMT3A*-R882) were included. Upon generation of a “cyt” file, data was log normalized. Dimensionality reduction was performed generating UMAPs. UMAPs were then divided into 50×50 squares and for each square, we calculated the number of WT and DNMT3A R882 mutant cells in each square. The ratio of the number of *DNMT3A*-R882 mutant cells/number of WT cells was calculated for each square and visualized on the UMAP. The MFI of individual cell surface markers was also visualised.

### Flow cytometry phenotyping analysis

Flow cytometry data of colonies derived from single HSC/MPPs from each individual was analysed in FlowJo^TM^ v. 10.8.1 Software (BD BioSciences) employing the manual gating strategy described in **Fig. S1B**. The counts, MFI of markers of interest, and percentage of parent populations for each population of interest were exported and analysed in R v4.2.3.

Colonies were categorised based on their mature cell composition. Colonies with more than 50 events in the Ery gate (GlyA+/CD45-) were classified as Ery-containing. Colonies with more than 50 events in either the Early Neut gate (CD56-/CD14-/CD15+/CD66b-), or in the Intermediate Neut gate (CD56-/CD14-/CD15+/CD66b+/CD11b-), or in the Mature Neut gate (CD56-/CD14-/CD15+/CD66b+/CD11b+) were classified as Neutrophil-containing. Monocyte-containing colonies were defined as colonies with either more than 400 cells in the Mono gate (CD14+/CD11b+/CD15-, CD14+/CD33+/CD15-, CD14+/CD11b+/CD66b-, based on staining panel employed) or between 150 and 400 cells in the Mono gate, provided this count was not lower than 2.5% of the total single cell number. These classification criteria are not mutually exclusive, so that a single colony could carry multiple labels, or none. Thresholds were established to effectively differentiate genuine cell populations from noise, following a comprehensive visual analysis of flow-cytometry data across all colonies. Data for all colonies analysed in this study is reported in **Table S2**.

To interrogate the effect of *DNMT3A* genotype (WT or R882) on each parameter of interest, we fitted linear mixed-effects models using the *lme4* v1.1-34 R package, with the restricted maximum likelihood (REML) default approach. *DNMT3A* genotype was included as fixed effect, while sample ID was modelled as random effect to account for inter-individual variability. P-value for fixed effect was computed using Satterthwaite’s method, as implemented in the *lmerTest* v3.1-3 package, and multiple testing correction was performed using the Benjamini-Hochberg method. Other statistical tests employed are listed in the relevant figure legends.

### WGS and phylogenetic tree generation and analysis

Reads were aligned to the human reference genome (GRCh37 – hg19) using BWA-MEM and CaVEMan and Pindels were employed to call indels and SNVs, respectively. Phylogenetic trees were constructed, and terminal branch length was corrected for sequencing depth within each individual colony, using the published approach as in ^1,108^. Shannon diversity index was employed as measure of clonality. To compare phylogenetic trees of different sizes, every tree (or every sub-section of interest, eg. CHIP1 *DNMT3A*-R882 mutant section) was downsampled to 100 randomly generated trees of 33 tips of size. CHIP1 WT section did not undergo downsampling, being it already 33 tips in size. Shannon diversity index was then calculated by summing the probability of each clade, defined by the number of tips in the clade over the total number of tips in the tree. Clades were defined as the clones present at 60 mutations of molecular time.

### Methylation analysis

The samples were processed by the in-house Sanger methylation sequencing pipeline which follows the bioinformatic steps detailed in version 1.4 of the nf-core methylseq pipeline (https://nf-co.re/methylseq/1.4/). A median whole genome depth of 1X or more was attained in 72 of the 96 samples, and 54 of these samples had a median autosomal CpG depth >=8 and were carried forward for further analysis. In total 27.6 million CpG sites had non-zero coverage. CpGs satisfying any of the following criteria were removed: i) Aggregate coverage that differed by a factor of 3 from the median coverage; ii) Aggregate forward and reverse strand coverage differing by more than a factor of 2, iii) >30% of the samples exhibiting CpG depth<8; iv) CpG overlapped a SNP found in the corresponding WGS samples. This left 22.8 million CpG sites for further analysis. Global DNA methylation levels were then calculated as mean CpG methylation across autosomes in myeloid colonies.

### scRNA-seq analysis

Analysis was performed with Python v. 3.8.10 and R v. 4.3.2. Reads were mapped to hg19-3.0.0 reference genome with cell ranger v. 6.0.1 using cell ranger count function options –-include-introns and –-force-cells, as recommended by 10X guidelines for neutrophils detection ^109^. Doublets were removed using scrublet v. 0.2.3. Quality control, log-normalisation, and highly variable gene selection was performed following the Scanpy (v 1.9.8) workflow and inbuilt package methods. For quality control, we excluded genes expressed in fewer than three cells and removed cells with less than 500 detected genes, with total counts below 13,000 or above 85,000, or mitochondrial gene content above 15%. Unwanted sources of variability, such as total counts and mitochondrial gene content were regressed out.

Batch correction across libraries was performed with the BBKNN algorithm ^110^ (package v.1.6.0). Following dimensionality reduction and UMAP visualisation, the data was clustered with the Leiden algorithm implemented in Scanpy. Clustering resolution and cluster annotation were guided by label transfer from publicly available reference datasets ^62,111,112^, using the Azimuth v. 0.4.6 or the symphony v 0.1.1 R packages, and by inspecting the top50 markers per cluster (**Table S3**). Diffusion pseudotime was performed employing the built in Scanpy function ^113^.

Differential gene expression (DGE) analysis was performed using edgeR (v4.0.16) after generating two pseudobulks per library. A DGE model was fit per cell type and treatment condition, controlling for differences in sort strategy per library (design matrix= 0∼genotype+sort) to evaluate the effect of the *DNMT3A*-R882 mutation in all tested conditions. An additional model was fit to evaluate the effect of interferon treatment independently of mutational status (design matrix= 0∼genotype+sort+treatment+genotype:treatment; differentially expressed genes obtained by extracting “treatment” coefficient). Differentially expressed genes were ranked by the product of the sign of the fold change with the –log10 of the Pvalue and gene set enrichment analysis (GSEA) was performed with *fgsea* v1.28.0 package on a combined collection of MSigDB Hallmark gene sets (v 2023.2, human) and gene sets of interest^60,61,82,83,114^ (**Table S3)**.

To identify transcriptional signatures defining *DNMT3A*-R882 mutant neutrophils at steady state and post IFNγ exposure, we selected genes with a false discovery rate (FDR) threshold of 0.05 and log fold change threshold greater than 1. Gene set variation analysis (GSVA v1.50.5) was performed to assess the enrichment of these signatures in publicly available datasets.

Inference of statistically significant cellular communication networks was performed with CellChat v2.1.2, employing the “Secreted Signaling” database included in the package.

### bulk RNA-seq analysis

Analysis was performed in R v 4.3.2. FASTQ files were trimmed for low quality end and adapter content using *trimgalore* v0.6.10 package (cutadapt v2.6, FastQC v0.12.1). Reads were aligned to GRCh37 reference genome with *star* v.2.0.7 package. Features counts were computed with *subreads* v2.0.8 (-t exome). QC was performed and samples with mitochondrial and ribosomal content under 5%, and genes with more than 10 counts in over 20% of samples were retained for downstream analysis. edgeR v4.0.16 package was used for TMM normalisation prior to principal component analysis (PCA), and for differential gene expression (DGE) analysis. A DGE model was fit per cell type and per condition to assess the effect on *DNMT3A* genotype in all tested contexts. Gene set enrichment analysis (GSEA) was performed with the *fgsea* v1.28.0 package on a combined collection of MSigDB Hallmark gene sets (v 2023.2, human) and gene sets of interest reported in **Table S4**.

### Statistical analyses and software

**<u>In addition to all software described in the specific analytic and statistical pipeline described above,</u>** flow cytometry data was acquired with FACSDiva v. 6.2 (BD Biosciences). Schematics of experiment design were created with BioRender.com and covered by proper licence (Agreement numbers: Fig 1: UA28UGI0ZN, Fig2: VF28UGI139, Fig3: NY28UGI14Z, Fig 4: XO28UGI1AF, Fig S3: AB28UGI18L).

## Acknowledgments

We would like to express our enormous gratitude for the generous donation of human tissue by the BM, PB and mPB donors. We would also like to thank the Cambridge NIHR BRC Cell Phenotyping Hub for their flow cytometry services and advice, the Jeffrey Cheah Biomedical Centre NGS Core Facility for the RNA-seq library preparation, and the CRUK Cambridge Institute genomics centre for sequencing. This research was supported by the CIMR Flow Cytometry Core Facility. We wish to thank Reiner Shulte and Gabriela Grondys-Kotarba for their advice and support in cell sorting. We thank Joanna Baxter, the team of the Cambridge Stem Cell Biobank and Evangelia Giannelou for their support in sample management. We would also like to acknowledge the Francis Crick core flow cytometry and advance sequencing facilities, as well as the Anthony Nolan charity for provision of cord blood for experiments performed at the Crick. Finally, we thank Stephanie Xie for critical reading of the manuscript.

## Funding

British Council BIRAX: 47BX16ELLS (EL, LS)

Wellcome – Royal Society Sir Henry Dale Fellowship 107630/Z/15/Z (EL)

Blood Cancer United (previously Leukemia and Lymphoma Society) 7035-24 (EL, GV)

Blood Cancer UK 24008 (EL)

Core support grants by Wellcome and Medical Research Council (MRC) to the Wellcome-MRC Cambridge Stem Cell Institute 203151/Z/16/Z (EL)

The Alborada Trust (EL, JN)

ISF-IPMP-Israel Precision Medicine Program 3165/19 (LS)

ISF-NSFC 2427/18 (LS)

ISF 1123/21 (LS)

Ernest and Bonnie Beutler Research Program of Excellence in Genomic Medicine (LS)

ERC Consolidator grant (UDEMC) (LS).

LS is an incumbent of the Ruth and Louis Leland career development chair, Applebaum Foundation, the Bolton Hope foundation and the Anthony Beck foundation.

The WBH Foundation (PC)

Gates Cambridge Trust PhD Scholarship, Application Number: 10350885 (AV) 2017-2022. Cancer Research UK Cambridge Cancer Centre PhD fellowship [CTRQQR-2021\100012] (GM, AS) Maria Sklodowska-Curie National Research Institute of Oncology, Gliwice Branch, Poland and Fundacja im. Jakuba Potockiego, Poland (AK).

Core support grants by Cancer Research UK (CC2027), The UK Medical Research Council (CC2027) and the Welcome Trust (CC20-27) to the Francis Crick Institute (DB).

Kay Kendall Leukaemia foundation KKL 1397 (HHE)

European Hematology Association Kick-off grant (HHE).

The collection of samples and data from the SardiNIA longitudinal cohort study was supported by the Intramural Research Program of the NIH, National Institute on Aging (NIA) of the National Institute of Health (NIH) with contracts N01-AG-1-2109 and HHSN271201100005C; and by the European Union’s Horizon 2020 Research and Innovation Programme under grant agreement 633964 (ImmunoAgeing). **This research was funded in part by the Wellcome Trust (107630/Z/15/Z). For the purpose of Open Access, the author has applied a CC BY public copyright licence to any Author Accepted Manuscript version arising from this submission**.

## Contributions

Conceptualization: EL, LS

Methodology: EL, LS, AV, GM, HHE, DB, AT

Investigation: AV, GM, HHE, AT, DH, AS, TB, YM, AD, NC

Visualization: GM, AV, DH, AK, KS

Formal analysis: GM, AV, DH, AK, KS, EM, JL, NW, WD, HHE

Data curation: GM, AV, HB, KS, DH, AK Validation: EL, GM, AV, HB

Resources: LS, MaM, AA, MC, JN, EF, VO, MiM, EMK, FC, PC, DB, GSV, MAF

Writing—original draft: EL, AV, GM

Writing—review & editing: EL, AV, GM

Supervision: EL, LS, GSV, JN, DB

Project management: EL, AV, GM

Funding acquisition: EL, LS, GSV, JN

GM and AV contributed equally and reserve the right to list their name first on their CV and professional applications.

## References

1. Mitchell, E., Spencer Chapman, M., Williams, N., Dawson, K.J., Mende, N., Calderbank, E.F., Jung, H., Mitchell, T., Coorens, T.H.H., Spencer, D.H., et al. (2022). Clonal dynamics of haematopoiesis across the human lifespan. Nature 606, 343–350. 10.1038/s41586-022-04786-y.

2. Jaiswal, S., Fontanillas, P., Flannick, J., Manning, A., Grauman, P.V., Mar, B.G., Lindsley, R.C., Mermel, C.H., Burtt, N., Chavez, A., et al. (2014). Age-Related Clonal Hematopoiesis Associated with Adverse Outcomes. N. Engl. J. Med. 371, 2488–2498. 10.1056/NEJMoa1408617.

3. Genovese, G., Kähler, A.K., Handsaker, R.E., Lindberg, J., Rose, S.A., Bakhoum, S.F., Chambert, K., Mick, E., Neale, B.M., Fromer, M., et al. (2014). Clonal Hematopoiesis and Blood-Cancer Risk Inferred from Blood DNA Sequence. N. Engl. J. Med. 371, 2477–2487. 10.1056/NEJMoa1409405.

4. Tall, A.R., and Fuster, J.J. (2022). Clonal hematopoiesis in cardiovascular disease and therapeutic implications. Nat. Cardiovasc. Res. 1, 116–124. 10.1038/s44161-021-00015-3.

5. Wong, W.J., Emdin, C., Bick, A.G., Zekavat, S.M., Niroula, A., Pirruccello, J.P., Dichtel, L., Griffin, G., Uddin, M.M., Gibson, C.J., et al. (2023). Clonal haematopoiesis and risk of chronic liver disease. Nature 616, 747–754. 10.1038/s41586-023-05857-4.

6. Agrawal, M., Niroula, A., Cunin, P., McConkey, M., Shkolnik, V., Kim, P.G., Wong, W.J., Weeks, L.D., Lin, A.E., Miller, P.G., et al. (2022). TET2-mutant clonal hematopoiesis and risk of gout. Blood 140, 1094–1103. 10.1182/blood.2022015384.

7. Kim, P.G., Niroula, A., Shkolnik, V., McConkey, M., Lin, A.E., Słabicki, M., Kemp, J.P., Bick, A., Gibson, C.J., Griffin, G., et al. (2021). Dnmt3a-mutated clonal hematopoiesis promotes osteoporosis. J. Exp. Med. 218, e20211872. 10.1084/jem.20211872.

8. Wang, H., Divaris, K., Pan, B., Li, X., Lim, J.-H., Saha, G., Barovic, M., Giannakou, D., Korostoff, J.M., Bing, Y., et al. (2024). Clonal hematopoiesis driven by mutated DNMT3A promotes inflammatory bone loss. Cell 187, 3690–3711.e19. 10.1016/j.cell.2024.05.003.

9. Miller, P.G., Qiao, D., Rojas-Quintero, J., Honigberg, M.C., Sperling, A.S., Gibson, C.J., Bick, A.G., Niroula, A., McConkey, M.E., Sandoval, B., et al. (2022). Association of clonal hematopoiesis with chronic obstructive pulmonary disease. Blood 139, 357–368. 10.1182/blood.2021013531.

10. Kessler, M.D., Damask, A., O’Keeffe, S., Banerjee, N., Li, D., Watanabe, K., Marketta, A., Van Meter, M., Semrau, S., Horowitz, J., et al. (2022). Common and rare variant associations with clonal haematopoiesis phenotypes. Nature 612, 301–309. 10.1038/s41586-022-05448-9.

11. Pich, O., Bernard, E., Zagorulya, M., Rowan, A., Pospori, C., Slama, R., Encabo, H.H., O’Sullivan, J., Papazoglou, D., Anastasiou, P., et al. (2025). Tumor-Infiltrating Clonal Hematopoiesis. N. Engl. J. Med. 392, 1594–1608. 10.1056/NEJMoa2413361.

12. Tian, R., Wiley, B., Liu, J., Zong, X., Truong, B., Zhao, S., Uddin, M.M., Niroula, A., Miller, C.A., Mukherjee, S., et al. (2023). Clonal Hematopoiesis and Risk of Incident Lung Cancer. J. Clin. Oncol. 41, 1423–1433. 10.1200/JCO.22.00857.

13. López-Otín, C., Blasco, M.A., Partridge, L., Serrano, M., and Kroemer, G. (2023). Hallmarks of aging: An expanding universe. Cell 186, 243–278. 10.1016/j.cell.2022.11.001.

14. Bogeska, R., Mikecin, A.-M., Kaschutnig, P., Fawaz, M., Büchler-Schäff, M., Le, D., Ganuza, M., Vollmer, A., Paffenholz, S.V., Asada, N., et al. (2022). Inflammatory exposure drives long-lived impairment of hematopoietic stem cell self-renewal activity and accelerated aging. Cell Stem Cell 29, 1273–1284.e8. 10.1016/j.stem.2022.06.012.

15. Hormaechea-Agulla, D., Matatall, K.A., Le, D.T., Kain, B., Long, X., Kus, P., Jaksik, R., Challen, G.A., Kimmel, M., and King, K.Y. (2021). Chronic infection drives Dnmt3a-loss-of-function clonal hematopoiesis via IFNγ signaling. Cell Stem Cell 28, 1428–1442.e6. 10.1016/j.stem.2021.03.002.

16. Trowbridge, J.J., and Starczynowski, D.T. (2021). Innate immune pathways and inflammation in hematopoietic aging, clonal hematopoiesis, and MDS. J. Exp. Med. 218. 10.1084/jem.20201544.

17. Pietras, E.M. (2017). Inflammation: a key regulator of hematopoietic stem cell fate in health and disease. Blood 130, 1693–1698. 10.1182/blood-2017-06-780882.

18. Zioni, N., Bercovich, A.A., Chapal-Ilani, N., Bacharach, T., Rappoport, N., Solomon, A., Avraham, R., Kopitman, E., Porat, Z., Sacma, M., et al. (2023). Inflammatory signals from fatty bone marrow support DNMT3A driven clonal hematopoiesis. Nat. Commun. 14, 2070. 10.1038/s41467-023-36906-1.

19. Xie, M., Lu, C., Wang, J., McLellan, M.D., Johnson, K.J., Wendl, M.C., McMichael, J.F., Schmidt, H.K., Yellapantula, V., Miller, C.A., et al. (2014). Age-related mutations associated with clonal hematopoietic expansion and malignancies. Nat. Med. 20, 1472– 1478. 10.1038/nm.3733.

20. McKerrell, T., Park, N., Moreno, T., Grove, C.S., Ponstingl, H., Stephens, J., Understanding Society Scientific Group, Crawley, C., Craig, J., Scott, M.A., et al. (2015). Leukemia-associated somatic mutations drive distinct patterns of age-related clonal hemopoiesis. Cell Rep. 10, 1239–1245. 10.1016/j.celrep.2015.02.005.

21. Thol, F., Damm, F., Lüdeking, A., Winschel, C., Wagner, K., Morgan, M., Yun, H., Göhring, G., Schlegelberger, B., Hoelzer, D., et al. (2011). Incidence and Prognostic Influence of DNMT3A Mutations in Acute Myeloid Leukemia. J. Clin. Oncol. 29, 2889–2896. 10.1200/JCO.2011.35.4894.

22. Brunetti, L., Gundry, M.C., and Goodell, M.A. (2017). DNMT3A in Leukemia. Cold Spring Harb. Perspect. Med. 7, a030320. 10.1101/cshperspect.a030320.

23. Russler-Germain, D.A., Spencer, D.H., Young, M.A., Lamprecht, T.L., Miller, C.A., Fulton, R., Meyer, M.R., Erdmann-Gilmore, P., Townsend, R.R., Wilson, R.K., et al. (2014). The R882H DNMT3A Mutation Associated with AML Dominantly Inhibits Wild-Type DNMT3A by Blocking Its Ability to Form Active Tetramers. Cancer Cell 25, 442–454. 10.1016/j.ccr.2014.02.010.

24. Ley, T.J., Ding, L., Walter, M.J., McLellan, M.D., Lamprecht, T., Larson, D.E., Kandoth, C., Payton, J.E., Baty, J., Welch, J., et al. (2010). DNMT3A Mutations in Acute Myeloid Leukemia. N. Engl. J. Med. 363, 2424–2433. 10.1056/NEJMoa1005143.

25. Abelson, S., Collord, G., Ng, S.W.K., Weissbrod, O., Mendelson Cohen, N., Niemeyer, E., Barda, N., Zuzarte, P.C., Heisler, L., Sundaravadanam, Y., et al. (2018). Prediction of acute myeloid leukaemia risk in healthy individuals. Nature 559, 400–404. 10.1038/s41586-018-0317-6.

26. Desai, P., Mencia-Trinchant, N., Savenkov, O., Simon, M.S., Cheang, G., Lee, S., Samuel, M., Ritchie, E.K., Guzman, M.L., Ballman, K.V., et al. (2018). Somatic mutations precede acute myeloid leukemia years before diagnosis. Nat. Med. 24, 1015–1023. 10.1038/s41591-018-0081-z.

27. Emperle, M., Rajavelu, A., Kunert, S., Arimondo, P.B., Reinhardt, R., Jurkowska, R.Z., and Jeltsch, A. (2018). The DNMT3A R882H mutant displays altered flanking sequence preferences. Nucleic Acids Res. 46, 3130–3139. 10.1093/nar/gky168.

28. Nam, A.S., Dusaj, N., Izzo, F., Murali, R., Myers, R.M., Mouhieddine, T.H., Sotelo, J., Benbarche, S., Waarts, M., Gaiti, F., et al. (2022). Single-cell multi-omics of human clonal hematopoiesis reveals that DNMT3A R882 mutations perturb early progenitor states through selective hypomethylation. Nat. Genet., 1–13. 10.1038/s41588-022-01179-9.

29. Koya, J., Kataoka, K., Sato, T., Bando, M., Kato, Y., Tsuruta-Kishino, T., Kobayashi, H., Narukawa, K., Miyoshi, H., Shirahige, K., et al. (2016). DNMT3A R882 mutants interact with polycomb proteins to block haematopoietic stem and leukaemic cell differentiation. Nat. Commun. 7, 10924. 10.1038/ncomms10924.

30. Fabre, M.A., de Almeida, J.G., Fiorillo, E., Mitchell, E., Damaskou, A., Rak, J., Orrù, V., Marongiu, M., Chapman, M.S., Vijayabaskar, M.S., et al. (2022). The longitudinal dynamics and natural history of clonal haematopoiesis. Nature 606, 335–342. 10.1038/s41586-022-04785-z.

31. Watson, C.J., Papula, A.L., Poon, G.Y.P., Wong, W.H., Young, A.L., Druley, T.E., Fisher, D.S., and Blundell, J.R. (2020). The evolutionary dynamics and fitness landscape of clonal hematopoiesis. Science 367, 1449–1454. 10.1126/science.aay9333.

32. Challen, G.A., Sun, D., Jeong, M., Luo, M., Jelinek, J., Berg, J.S., Bock, C., Vasanthakumar, A., Gu, H., Xi, Y., et al. (2012). Dnmt3a is essential for hematopoietic stem cell differentiation. Nat. Genet. 44, 23. 10.1038/ng.1009.

33. Jeong, M., Park, H.J., Celik, H., Ostrander, E.L., Reyes, J.M., Guzman, A., Rodriguez, B., Lei, Y., Lee, Y., Ding, L., et al. (2018). Loss of Dnmt3a Immortalizes Hematopoietic Stem Cells In Vivo. Cell Rep. 23, 1–10. 10.1016/j.celrep.2018.03.025.

34. Guryanova, O.A., Lieu, Y.K., Garrett-Bakelman, F.E., Spitzer, B., Glass, J.L., Shank, K., Martinez, A.B.V., Rivera, S.A., Durham, B.H., Rapaport, F., et al. (2016). *Dnmt3a* regulates myeloproliferation and liver-specific expansion of hematopoietic stem and progenitor cells. Leukemia 30, 1133. 10.1038/leu.2015.358.

35. Guryanova, O.A., Shank, K., Spitzer, B., Luciani, L., Koche, R.P., Garrett-Bakelman, F.E., Ganzel, C., Durham, B.H., Mohanty, A., Hoermann, G., et al. (2016). *DNMT3A* mutations promote anthracycline resistance in acute myeloid leukemia via impaired nucleosome remodeling. Nat. Med. 22, 1488. 10.1038/nm.4210.

36. Shlush, L.I., Zandi, S., Mitchell, A., Chen, W.C., Brandwein, J.M., Gupta, V., Kennedy, J.A., Schimmer, A.D., Schuh, A.C., Yee, K.W., et al. (2014). Identification of pre-leukaemic haematopoietic stem cells in acute leukaemia. Nature 506, 328–333. 10.1038/nature13038.

37. Karpova, D., Huerga Encabo, H., Donato, E., Calderazzo, S., Scherer, M., Llorian-Sopena, M., Leppä, A.-M., Würth, R., Stelmach, P., Papazoglou, D., et al. (2025). Clonal hematopoiesis landscape in frequent blood donors. Blood 145, 2411–2423. 10.1182/blood.2024027999.

38. Smith, A.M., Verdoni, A.M., Abel, H.J., Chen, D.Y., Ketkar, S., Leight, E.R., Miller, C.A., and Ley, T.J. (2022). Somatic Dnmt3a inactivation leads to slow, canonical DNA methylation loss in murine hematopoietic cells. iScience 25. 10.1016/j.isci.2022.104004.

39. Ramabadran, R., Wang, J.H., Reyes, J.M., Guzman, A.G., Gupta, S., Rosas, C., Brunetti, L., Gundry, M.C., Tovy, A., Long, H., et al. (2023). DNMT3A-coordinated splicing governs the stem state switch towards differentiation in embryonic and haematopoietic stem cells. Nat. Cell Biol., 1–12. 10.1038/s41556-023-01109-9.

40. Izzo, F., Lee, S.C., Poran, A., Chaligne, R., Gaiti, F., Gross, B., Murali, R.R., Deochand, S.D., Ang, C., Jones, P.W., et al. (2020). DNA methylation disruption reshapes the hematopoietic differentiation landscape. Nat. Genet., 1–10. 10.1038/s41588-020-0595-4.

41. Isobe, T., Kucinski, I., Barile, M., Wang, X., Hannah, R., Bastos, H.P., Chabra, S., Vijayabaskar, M.S., Sturgess, K.H.M., Williams, M.J., et al. (2023). Preleukemic single-cell landscapes reveal mutation-specific mechanisms and gene programs predictive of AML patient outcomes. Cell Genomics, 100426. 10.1016/j.xgen.2023.100426.

42. Jakobsen, N.A., Turkalj, S., Zeng, A.G.X., Stoilova, B., Metzner, M., Rahmig, S., Nagree, M.S., Shah, S., Moore, R., Usukhbayar, B., et al. (2024). Selective advantage of mutant stem cells in human clonal hematopoiesis is associated with attenuated response to inflammation and aging. Cell Stem Cell 31, 1127–1144.e17. 10.1016/j.stem.2024.05.010.

43. Furer, N., Rappoport, N., Milman, O., Tavor, S., Lifshitz, A., Bercovich, A., Ben-Kiki, O., Danin, A., Kedmi, M., Shipony, Z., et al. (2025). A reference model of circulating hematopoietic stem cells across the lifespan with applications to diagnostics. Nat. Med., 1–10. 10.1038/s41591-025-03716-5.

44. SanMiguel, J.M., Eudy, E., Loberg, M.A., Young, K.A., Mistry, J.J., Mujica, K.D., Schwartz, L.S., Stearns, T.M., Challen, G.A., and Trowbridge, J.J. (2022). Distinct Tumor Necrosis Factor Alpha Receptors Dictate Stem Cell Fitness versus Lineage Output in Dnmt3a-Mutant Clonal Hematopoiesis. Cancer Discov. 12, 2763–2773. 10.1158/2159-8290.CD-22-0086.

45. Liao, M., Chen, R., Yang, Y., He, H., Xu, L., Jiang, Y., Guo, Z., He, W., Jiang, H., and Wang, J. (2022). Aging-elevated inflammation promotes DNMT3A R878H-driven clonal hematopoiesis. Acta Pharm. Sin. B 12, 678–691. 10.1016/j.apsb.2021.09.015.

46. Rodrigues, K.B., Gopakumar, J., Weng, Z., Mitchell, S.R., Maurer, M., Nachun, D., Eulalio, T., Estrada, D., Mazumder, T., Ma, L., et al. (2023). Multi-Omic Profiling of Macrophages Lacking Tet2 or Dnmt3a Reveals Mechanisms of Hyper-Inflammation in Clonal Hematopoiesis. Blood 142, 1163. 10.1182/blood-2023-187890.

47. Rauch, P.J., Gopakumar, J., Silver, A.J., Nachun, D., Ahmad, H., McConkey, M., Nakao, T., Bosse, M., Rentz, T., Vivanco Gonzalez, N., et al. (2023). Loss-of-function mutations in Dnmt3a and Tet2 lead to accelerated atherosclerosis and concordant macrophage phenotypes. Nat. Cardiovasc. Res. 2, 805–818. 10.1038/s44161-023-00326-7.

48. Cobo, I., Tanaka, T.N., Chandra Mangalhara, K., Lana, A., Yeang, C., Han, C., Schlachetzki, J., Challcombe, J., Fixsen, B.R., Sakai, M., et al. (2022). DNA methyltransferase 3 alpha and TET methylcytosine dioxygenase 2 restrain mitochondrial DNA-mediated interferon signaling in macrophages. Immunity 55, 1386–1401.e10. 10.1016/j.immuni.2022.06.022.

49. Belluschi, S., Calderbank, E.F., Ciaurro, V., Pijuan-Sala, B., Santoro, A., Mende, N., Diamanti, E., Sham, K.Y.C., Wang, X., Lau, W.W.Y., et al. (2018). Myelo-lymphoid lineage restriction occurs in the human haematopoietic stem cell compartment before lymphoid-primed multipotent progenitors. Nat. Commun. 9, 4100. 10.1038/s41467-018-06442-4.

50. Vaisvila, R., Ponnaluri, V.K.C., Sun, Z., Langhorst, B.W., Saleh, L., Guan, S., Dai, N., Campbell, M.A., Sexton, B.S., Marks, K., et al. (2021). Enzymatic methyl sequencing detects DNA methylation at single-base resolution from picograms of DNA. Genome Res. 31, 1280–1289. 10.1101/gr.266551.120.

51. Lee-Six, H., Øbro, N.F., Shepherd, M.S., Grossmann, S., Dawson, K., Belmonte, M., Osborne, R.J., Huntly, B.J.P., Martincorena, I., Anderson, E., et al. (2018). Population dynamics of normal human blood inferred from somatic mutations. Nature 561, 473–478. 10.1038/s41586-018-0497-0.

52. Notta, F., Doulatov, S., Laurenti, E., Poeppl, A., Jurisica, I., and Dick, J.E. (2011). Isolation of Single Human Hematopoietic Stem Cells Capable of Long-Term Multilineage Engraftment. Science 333, 218–221. 10.1126/science.1201219.

53. Jamieson, C.H.M., Gotlib, J., Durocher, J.A., Chao, M.P., Mariappan, M.R., Lay, M., Jones, C., Zehnder, J.L., Lilleberg, S.L., and Weissman, I.L. (2006). The JAK2 V617F mutation occurs in hematopoietic stem cells in polycythemia vera and predisposes toward erythroid differentiation. Proc. Natl. Acad. Sci. U. S. A. 103, 6224–6229. 10.1073/pnas.0601462103.

54. Tong, J., Sun, T., Ma, S., Zhao, Y., Ju, M., Gao, Y., Zhu, P., Tan, P., Fu, R., Zhang, A., et al. (2021). Hematopoietic Stem Cell Heterogeneity Is Linked to the Initiation and Therapeutic Response of Myeloproliferative Neoplasms. Cell Stem Cell 28, 502–513.e6. 10.1016/j.stem.2021.01.018.

55. Orrù, V., Steri, M., Sole, G., Sidore, C., Virdis, F., Dei, M., Lai, S., Zoledziewska, M., Busonero, F., Mulas, A., et al. (2013). Genetic Variants Regulating Immune Cell Levels in Health and Disease. Cell 155, 242–256. 10.1016/j.cell.2013.08.041.

56. Dai, Y., Xu, A., Li, J., Wu, L., Yu, S., Chen, J., Zhao, W., Sun, X.-J., and Huang, J. (2021). CytoTree: an R/Bioconductor package for analysis and visualization of flow and mass cytometry data. BMC Bioinformatics 22, 138. 10.1186/s12859-021-04054-2.

57. Evrard, M., Kwok, I.W.H., Chong, S.Z., Teng, K.W.W., Becht, E., Chen, J., Sieow, J.L., Penny, H.L., Ching, G.C., Devi, S., et al. (2018). Developmental Analysis of Bone Marrow Neutrophils Reveals Populations Specialized in Expansion, Trafficking, and Effector Functions. Immunity 48, 364–379.e8. 10.1016/j.immuni.2018.02.002.

58. Calzetti, F., Finotti, G., Tamassia, N., Bianchetto-Aguilera, F., Castellucci, M., Canè, S., Lonardi, S., Cavallini, C., Matte, A., Gasperini, S., et al. (2022). CD66b−CD64dimCD115− cells in the human bone marrow represent neutrophil-committed progenitors. Nat. Immunol. 23, 679–691. 10.1038/s41590-022-01189-z.

59. Encabo, H.H., Aramburu, I.V., Garcia-Albornoz, M., Piganeau, M., Wood, H., Song, A., Ferrelli, A., Sharma, A., Minutti, C.M., Domart, M.-C., et al. (2023). Loss of TET2 in human hematopoietic stem cells alters the development and function of neutrophils. Cell Stem Cell 30, 781–799.e9. 10.1016/j.stem.2023.05.004.

60. Montaldo, E., Lusito, E., Bianchessi, V., Caronni, N., Scala, S., Basso-Ricci, L., Cantaffa, C., Masserdotti, A., Barilaro, M., Barresi, S., et al. (2022). Cellular and transcriptional dynamics of human neutrophils at steady state and upon stress. Nat. Immunol., 1–14. 10.1038/s41590-022-01311-1.

61. Kirchberger, S., Shoeb, M.R., Lazic, D., Wenninger-Weinzierl, A., Fischer, K., Shaw, L.E., Nogueira, F., Rifatbegovic, F., Bozsaky, E., Ladenstein, R., et al. (2024). Comparative transcriptomics coupled to developmental grading via transgenic zebrafish reporter strains identifies conserved features in neutrophil maturation. Nat. Commun. 15, 1792. 10.1038/s41467-024-45802-1.

62. Kwok, A.J., Allcock, A., Ferreira, R.C., Cano-Gamez, E., Smee, M., Burnham, K.L., Zurke, Y.-X., McKechnie, S., Mentzer, A.J., Monaco, C., et al. (2023). Neutrophils and emergency granulopoiesis drive immune suppression and an extreme response endotype during sepsis. Nat. Immunol., 1–13. 10.1038/s41590-023-01490-5.

63. Lu, R., Wang, P., Parton, T., Zhou, Y., Chrysovergis, K., Rockowitz, S., Chen, W.-Y., Abdel-Wahab, O., Wade, P.A., Zheng, D., et al. (2016). Epigenetic Perturbations by Arg882-Mutated DNMT3A Potentiate Aberrant Stem Cell Gene-Expression Program and Acute Leukemia Development. Cancer Cell 30, 92–107. 10.1016/j.ccell.2016.05.008.

64. Khan, I., Amin, M.A., Eklund, E.A., and Gartel, A.L. (2024). Regulation of HOX gene expression in AML. Blood Cancer J. 14, 42. 10.1038/s41408-024-01004-y.

65. Huang, Z., Wei, J., Pan, X., Chen, X., and Lu, Z. (2024). A novel risk score model of lactate metabolism for predicting outcomes and immune signatures in acute myeloid leukemia. Sci. Rep. 14, 25742. 10.1038/s41598-024-76919-4.

66. Park, M.D., Le Berichel, J., Hamon, P., Wilk, C.M., Belabed, M., Yatim, N., Saffon, A., Boumelha, J., Falcomatà, C., Tepper, A., et al. (2024). Hematopoietic aging promotes cancer by fueling IL-1⍺–driven emergency myelopoiesis. Science 0, eadn0327. 10.1126/science.adn0327.

67. Caiado, F., and Manz, M.G. (2024). IL-1 in aging and pathologies of hematopoietic stem cells. Blood 144, 368–377. 10.1182/blood.2023023105.

68. Heimlich, J.B., Raddatz, M.A., Wells, J., Vlasschaert, C., Olson, S., Threadcraft, M., Foster, K., Boateng, E., Umbarger, K., Su, Y.R., et al. (2024). Invasive Assessment of Coronary Artery Disease in Clonal Hematopoiesis of Indeterminate Potential. Circ. Genomic Precis. Med. 17, e004415. 10.1161/CIRCGEN.123.004415.

69. Jaiswal, S., Natarajan, P., Silver, A.J., Gibson, C.J., Bick, A.G., Shvartz, E., McConkey, M., Gupta, N., Gabriel, S., Ardissino, D., et al. (2017). Clonal Hematopoiesis and Risk of Atherosclerotic Cardiovascular Disease. 10.1056/NEJMoa1701719. 10.1056/NEJMoa1701719.

70. Pekayvaz, K., Losert, C., Knottenberg, V., Gold, C., van Blokland, I.V., Oelen, R., Groot, H.E., Benjamins, J.W., Brambs, S., Kaiser, R., et al. (2024). Multiomic analyses uncover immunological signatures in acute and chronic coronary syndromes. Nat. Med. 30, 1696– 1710. 10.1038/s41591-024-02953-4.

71. Higazi, A.A.-R., Lavi, E., Bdeir, K., Ulrich, A.M., Jamieson, D.G., Rader, D.J., Usher, D.C., Kane, W., Ganz, T., and Cines, D.B. (1997). Defensin Stimulates the Binding of Lipoprotein (a) to Human Vascular Endothelial and Smooth Muscle Cells. Blood 89, 4290–4298. 10.1182/blood.V89.12.4290.

72. Abu-Fanne, R., Maraga, E., Abd-Elrahman, I., Hankin, A., Blum, G., Abdeen, S., Hijazi, N., Cines, D.B., and Higazi, A.A.-R. (2016). α-Defensins Induce a Post-translational Modification of Low Density Lipoprotein (LDL) That Promotes Atherosclerosis at Normal Levels of Plasma Cholesterol*. J. Biol. Chem. 291, 2777–2786. 10.1074/jbc.M115.669812.

73. Quinn, K.L., Henriques, M., Tabuchi, A., Han, B., Yang, H., Cheng, W.-E., Tole, S., Yu, H., Luo, A., Charbonney, E., et al. (2011). Human Neutrophil Peptides Mediate Endothelial-Monocyte Interaction, Foam Cell Formation, and Platelet Activation. Arterioscler. Thromb. Vasc. Biol. 31, 2070–2079. 10.1161/ATVBAHA.111.227116.

74. Jin, S., Plikus, M.V., and Nie, Q. (2025). CellChat for systematic analysis of cell–cell communication from single-cell transcriptomics. Nat. Protoc. 20, 180–219. 10.1038/s41596-024-01045-4.

75. Cersosimo, F., Lonardi, S., Ulivieri, C., Martini, P., Morrione, A., Vermi, W., Giordano, A., and Giurisato, E. (2024). CSF-1R in Cancer: More than a Myeloid Cell Receptor. Cancers 16, 282. 10.3390/cancers16020282.

76. Wang, Q., Lu, Y., Li, R., Jiang, Y., Zheng, Y., Qian, J., Bi, E., Zheng, C., Hou, J., Wang, S., et al. (2018). Therapeutic effects of CSF1R-blocking antibodies in multiple myeloma. Leukemia 32, 176–183. 10.1038/leu.2017.193.

77. Edwards, D.K., V., Watanabe-Smith, K., Rofelty, A., Damnernsawad, A., Laderas, T., Lamble, A., Lind, E.F., Kaempf, A., Mori, M., Rosenberg, M., et al. (2019). CSF1R inhibitors exhibit antitumor activity in acute myeloid leukemia by blocking paracrine signals from support cells. Blood 133, 588–599. 10.1182/blood-2018-03-838946.

78. Terabe, M., Matsui, S., Noben-Trauth, N., Chen, H., Watson, C., Donaldson, D.D., Carbone, D.P., Paul, W.E., and Berzofsky, J.A. (2000). NKT cell–mediated repression of tumor immunosurveillance by IL-13 and the IL-4R–STAT6 pathway. Nat. Immunol. 1, 515–520. 10.1038/82771.

79. Cai, X., Bowman, R.L., and Trowbridge, J.J. (2025). Clonal hematopoiesis in myeloid malignancies and solid tumors. Nat. Cancer, 1–12. 10.1038/s43018-025-01014-0.

80. Kleppe, M., Comen, E., Wen, H.Y., Bastian, L., Blum, B., Rapaport, F.T., Keller, M., Granot, Z., Socci, N., Viale, A., et al. (2015). Somatic mutations in leukocytes infiltrating primary breast cancers. Npj Breast Cancer 1, 15005. 10.1038/npjbcancer.2015.5.

81. Akkari, L., Amit, I., Bronte, V., Fridlender, Z.G., Gabrilovich, D.I., Ginhoux, F., Hedrick, C.C., and Ostrand-Rosenberg, S. (2024). Defining myeloid-derived suppressor cells. Nat. Rev. Immunol. 24, 850–857. 10.1038/s41577-024-01062-0.

82. Alshetaiwi, H., Pervolarakis, N., McIntyre, L.L., Ma, D., Nguyen, Q., Rath, J.A., Nee, K., Hernandez, G., Evans, K., Torosian, L., et al. (2020). Defining the emergence of myeloid-derived suppressor cells in breast cancer using single-cell transcriptomics. Sci. Immunol. 5, eaay6017. 10.1126/sciimmunol.aay6017.

83. van Galen, P., Hovestadt, V., Wadsworth II, M.H., Hughes, T.K., Griffin, G.K., Battaglia, S., Verga, J.A., Stephansky, J., Pastika, T.J., Lombardi Story, J., et al. (2019). Single-Cell RNA-Seq Reveals AML Hierarchies Relevant to Disease Progression and Immunity. Cell 176, 1265–1281.e24. 10.1016/j.cell.2019.01.031.

84. Watson, C.J., Poon, G.Y.P., MacGregor, H.A.J., Fonseca, A.V.A., Apostolidou, S., Gentry-Maharaj, A., Menon, U., and Blundell, J.R. (2024). Evolutionary dynamics in the decades preceding acute myeloid leukaemia. Preprint at bioRxiv, 10.1101/2024.07.05.602251 10.1101/2024.07.05.602251.

85. Braun, T.P., Estabrook, J., Schonrock, Z., Curtiss, B.M., Darmusey, L., Macaraeg, J., Enright, T., Coblentz, C., Callahan, R., Yashar, W., et al. (2023). Asxl1 deletion disrupts MYC and RNA polymerase II function in granulocyte progenitors. Leukemia 37, 478–487. 10.1038/s41375-022-01792-x.

86. Tovy, A., Rosas, C., Gaikwad, A.S., Medrano, G., Zhang, L., Reyes, J.M., Huang, Y.-H., Arakawa, T., Kurtz, K., Conneely, S.E., et al. (2020). Perturbed hematopoiesis in individuals with germline DNMT3A overgrowth Tatton-Brown-Rahman syndrome. Haematologica. 10.3324/haematol.2021.278990.

87. Tovy, A., Reyes, J.M., Gundry, M.C., Brunetti, L., Lee-Six, H., Petljak, M., Park, H.J., Guzman, A.G., Rosas, C., Jeffries, A.R., et al. (2020). Tissue-Biased Expansion of DNMT3A-Mutant Clones in a Mosaic Individual Is Associated with Conserved Epigenetic Erosion. Cell Stem Cell 27, 326–335.e4. 10.1016/j.stem.2020.06.018.

88. Gu, M., Kovilakam, S.C., Dunn, W.G., Marando, L., Barcena, C., Mohorianu, I., Smith, A., Kar, S.P., Fabre, M.A., Gerstung, M., et al. (2023). Multiparameter prediction of myeloid neoplasia risk. Nat. Genet. 55, 1523–1530. 10.1038/s41588-023-01472-1.

89. Avagyan, S., Henninger, J.E., Mannherz, W.P., Mistry, M., Yoon, J., Yang, S., Weber, M.C., Moore, J.L., and Zon, L.I. (2021). Resistance to inflammation underlies enhanced fitness in clonal hematopoiesis. Science 374, 768–772. 10.1126/science.aba9304.

90. Tran, B.T., Jeyanathan, V., Cao, R., Kaufmann, E., and King, K.Y. (2025). Hematopoietic stem and progenitor cells as a reservoir for trained immunity. eLife 14, e106610. 10.7554/eLife.106610.

91. Cheong, J.-G., Ravishankar, A., Sharma, S., Parkhurst, C.N., Grassmann, S.A., Wingert, C.K., Laurent, P., Ma, S., Paddock, L., Miranda, I.C., et al. (2023). Epigenetic memory of coronavirus infection in innate immune cells and their progenitors. Cell 186, 3882–3902.e24. 10.1016/j.cell.2023.07.019.

92. Li, X., Wang, H., Yu, X., Saha, G., Kalafati, L., Ioannidis, C., Mitroulis, I., Netea, M.G., Chavakis, T., and Hajishengallis, G. (2022). Maladaptive innate immune training of myelopoiesis links inflammatory comorbidities. Cell 185, 1709–1727.e18. 10.1016/j.cell.2022.03.043.

93. Christ, A., Günther, P., Lauterbach, M.A.R., Duewell, P., Biswas, D., Pelka, K., Scholz, C.J., Oosting, M., Haendler, K., Baßler, K., et al. (2018). Western Diet Triggers NLRP3-Dependent Innate Immune Reprogramming. Cell 172, 162–175.e14. 10.1016/j.cell.2017.12.013.

94. Bera, R., Chiu, M.-C., Huang, Y.-J., Liang, D.-C., Lee, Y.-S., and Shih, L.-Y. (2018). Genetic and Epigenetic Perturbations by DNMT3A-R882 Mutants Impaired Apoptosis through Augmentation of PRDX2 in Myeloid Leukemia Cells. Neoplasia 20, 1106–1120. 10.1016/j.neo.2018.08.013.

95. Yuan, X.-Q., Chen, P., Du, Y.-X., Zhu, K.-W., Zhang, D.-Y., Yan, H., Liu, H., Liu, Y.-L., Cao, S., Zhou, G., et al. (2019). Influence of DNMT3A R882 mutations on AML prognosis determined by the allele ratio in Chinese patients. J. Transl. Med. 17, 220. 10.1186/s12967-019-1959-3.

96. Huerga Encabo, H., Ulferts, R., Sharma, A., Beale, R., and Bonnet, D. (2021). Infecting human hematopoietic stem and progenitor cells with SARS-CoV-2. STAR Protoc. 2, 100903. 10.1016/j.xpro.2021.100903.

97. Tuval, A., Brilon, Y., Azogy, H., Moskovitz, Y., Leshkowitz, D., Salame, T.M., Minden, M.D., Tal, P., Rotter, V., Oren, M., et al. (2020). Pseudo-mutant P53 is a unique phenotype of *DNMT3A*-mutated pre-leukemia. Haematologica. 10.3324/haematol.2021.280329.

98. Acuna-Hidalgo, R., Sengul, H., Steehouwer, M., van de Vorst, M., Vermeulen, S.H., Kiemeney, L.A.L.M., Veltman, J.A., Gilissen, C., and Hoischen, A. (2017). Ultra-sensitive Sequencing Identifies High Prevalence of Clonal Hematopoiesis-Associated Mutations throughout Adult Life. Am. J. Hum. Genet. 101, 50–64. 10.1016/j.ajhg.2017.05.013.

99. Hiatt, J.B., Pritchard, C.C., Salipante, S.J., O’Roak, B.J., and Shendure, J. (2013). Single molecule molecular inversion probes for targeted, high-accuracy detection of low-frequency variation. Genome Res. 23, 843–854. 10.1101/gr.147686.112.

100. Mancini, M., Hasan, S.K., Ottone, T., Lavorgna, S., Ciardi, C., Angelini, D.F., Agostini, F., Venditti, A., and Lo-Coco, F. (2015). Two Novel Methods for Rapid Detection and Quantification of *DNMT3A* R882 Mutations in Acute Myeloid Leukemia. J. Mol. Diagn. 17, 179–184. 10.1016/j.jmoldx.2014.10.003.

101. Moore, L., Leongamornlert, D., Coorens, T.H.H., Sanders, M.A., Ellis, P., Dentro, S.C., Dawson, K.J., Butler, T., Rahbari, R., Mitchell, T.J., et al. (2020). The mutational landscape of normal human endometrial epithelium. Nature 580, 640–646. 10.1038/s41586-020-2214-z.

102. Bushnell, B., Rood, J., and Singer, E. (2017). BBMerge – Accurate paired shotgun read merging via overlap. PloS One 12, e0185056. 10.1371/journal.pone.0185056.

103. Martin, M. (2011). Cutadapt removes adapter sequences from high-throughput sequencing reads. EMBnet.journal 17, 10–12. 10.14806/ej.17.1.200.

104. Li, H. (2013). Aligning sequence reads, clone sequences and assembly contigs with BWA-MEM. Preprint at arXiv, 10.48550/arXiv.1303.3997 10.48550/arXiv.1303.3997.

105. McKenna, A., Hanna, M., Banks, E., Sivachenko, A., Cibulskis, K., Kernytsky, A., Garimella, K., Altshuler, D., Gabriel, S., Daly, M., et al. (2010). The Genome Analysis Toolkit: a MapReduce framework for analyzing next-generation DNA sequencing data. Genome Res. 20, 1297–1303. 10.1101/gr.107524.110.

106. Koboldt, D.C., Zhang, Q., Larson, D.E., Shen, D., McLellan, M.D., Lin, L., Miller, C.A., Mardis, E.R., Ding, L., and Wilson, R.K. (2012). VarScan 2: somatic mutation and copy number alteration discovery in cancer by exome sequencing. Genome Res. 22, 568–576. 10.1101/gr.129684.111.

107. Rimmer, A., Phan, H., Mathieson, I., Iqbal, Z., Twigg, S.R.F., WGS500 Consortium, Wilkie, A.O.M., McVean, G., and Lunter, G. (2014). Integrating mapping-, assembly– and haplotype-based approaches for calling variants in clinical sequencing applications. Nat. Genet. 46, 912–918. 10.1038/ng.3036.

108. Spencer Chapman, M., Wilk, C.M., Boettcher, S., Mitchell, E., Dawson, K., Williams, N., Müller, J., Kovtonyuk, L., Jung, H., Caiado, F., et al. (2024). Clonal dynamics after allogeneic haematopoietic cell transplantation. Nature, 1–9. 10.1038/s41586-024-08128-y.

109. Capturing Neutrophils in 10x Single Cell Gene Expression Data 10x Genomics. https://www.10xgenomics.com/support/software/cell-ranger/latest/tutorials/cr-tutorial-neutrophils.

110. Polański, K., Young, M.D., Miao, Z., Meyer, K.B., Teichmann, S.A., and Park, J.-E. (2020). BBKNN: fast batch alignment of single cell transcriptomes. Bioinforma. Oxf. Engl. 36, 964–965. 10.1093/bioinformatics/btz625.

111. Hao, Y., Hao, S., Andersen-Nissen, E., Mauck, W.M., Zheng, S., Butler, A., Lee, M.J., Wilk, A.J., Darby, C., Zager, M., et al. (2021). Integrated analysis of multimodal single-cell data. Cell 184, 3573–3587.e29. 10.1016/j.cell.2021.04.048.

112. Zeng, A.G.X., Iacobucci, I., Shah, S., Mitchell, A., Wong, G., Bansal, S., Chen, D., Gao, Q., Kim, H., Kennedy, J.A., et al. (2025). Single-cell Transcriptional Atlas of Human Hematopoiesis Reveals Genetic and Hierarchy-Based Determinants of Aberrant AML Differentiation. Blood Cancer Discov. 6, 307–324. 10.1158/2643-3230.BCD-24-0342.

113. Haghverdi, L., Büttner, M., Wolf, F.A., Buettner, F., and Theis, F.J. (2016). Diffusion pseudotime robustly reconstructs lineage branching. Nat. Methods 13, 845–848. 10.1038/nmeth.3971.

114. Hackert, N.S., Radtke, F.A., Exner, T., Lorenz, H.-M., Müller-Tidow, C., Nigrovic, P.A., Wabnitz, G., and Grieshaber-Bouyer, R. (2023). Human and mouse neutrophils share core transcriptional programs in both homeostatic and inflamed contexts. Nat. Commun. 14, 8133. 10.1038/s41467-023-43573-9.

