## Supplemental Tables and Figures for "*DNMT3A*-R882 mutations intrinsically drive dysfunctional neutropoiesis from human haematopoietic stem cells"

This supplemental material file contains:
- Tables S1-S4
- Figures S1-S4

**Table S1: summary of primary samples used in this study**
VAF = variant allele frequency; AML = Acute Myeloid Leukaemia; CHIP = Clonal Haematopoiesis of Indeterminate Potential; WT = wild type; PB = peripheral blood, mPB = mobilised PB; WGS = Whole Genome Sequencing, EM-seq = Enzymatic Methyl Sequencing, scRNA-seq= single cell RNA-seq. Of note, index sorting data is available for all samples for which colony phenotyping was performed. *CHIP2 was cultured in semisolid conditions (Methocult 4034, Stem Cell Technologies; see Methods).

| Donor | Diagnosis | Age | Gender | *DNMT3A R882* | Tissue | VAF colonies | VAF MNC | Colony pheno-typing | WGS | EM-seq | scRNA-seq |
|---|---|---|---|---|---|---|---|---|---|---|---|
| CHIP1 | CHIP | 63 | F | H | mPB | 0.340 | 0.340 | X | X | X | |
| CHIP2* | CHIP | 91 | M | H | PB | 0.451 | N/A | | X | | |
| CHIP3 | CHIP | 58 | F | H | PB | 0.490 | 0.69 | | | | |
| CHIP4 | CHIP | 80 | M | C | BM | 0.451 | 0.472 | X | | | |
| CHIP5 | CHIP | 41 | F | C | PB | 0.151 | 0.120 | | | | X |
| CHIP6 | CHIP | 81 | M | H | PB | 0.000 | 0.040 | | | | |
| AML1 | AML | 38 | M | H | PB | 0.268 | 0.200 | X | | | |
| AML2 | AML | 78 | M | H | PB | 0.372 | 0.480 | X | | | |
| AML3 | AML | 70 | M | C | PB | 0.239 | 0.475 | X | | | |
| AML4 | AML | 63 | F | C | PB | 0.375 | 0.447 | X | | | |
| AML5 | AML | 62 | M | H | PB | 0.393 | 0.484 | | | | |

**Table S2: summary of genotypes and phenotype from single HSC/MPP derived colonies**
**Tab 1:** variant calling results for CHIP1, CHIP3, CHIP4, CHIP6, and AML1-5, index sorting data and phenotyping data for CHIP1, CHIP4, AML1-4
**Tab 2:** number of colonies retrieved and genotyped for each donor.

**Table S3: scRNA-seq of myeloid cells derived from single *DNMT3A*-R882 or WT HSC/MPPs**
Tab 1: Top 50 marker genes of each of the clusters
Tab 2: differential gene expression analysis results for MUT vs WT untreated
Tab 3: differential gene expression analysis results for MUT vs WT IFNγ treated
Tab 4: differential gene expression analysis results for IFNγ vs untreated
Tab 5: signatures used for GSEA and GSVA analyses

**Table S4: editing efficiency and bulk RNA-seq of mice engrafted with edited *DNMT3A*-R882 HSPCs.**
Tab1: samples overview and VAF of genomic edits at the *DNMT3A* genomic locus at the time of bone marrow harvesting.
Tab2: differential gene expression analysis result for MUT vs WT untreated neutrophils
Tab3: differential gene expression analysis result for MUT vs WT IFNγ-treated neutrophils
Tab4: differential gene expression analysis result for MUT vs WT untreated neutrophil precursors
Tab5: differential gene expression analysis result for MUT vs WT IFNγ-treated neutrophil precursors
Tab6: signatures used for GSEA analysis

**Figure S1**

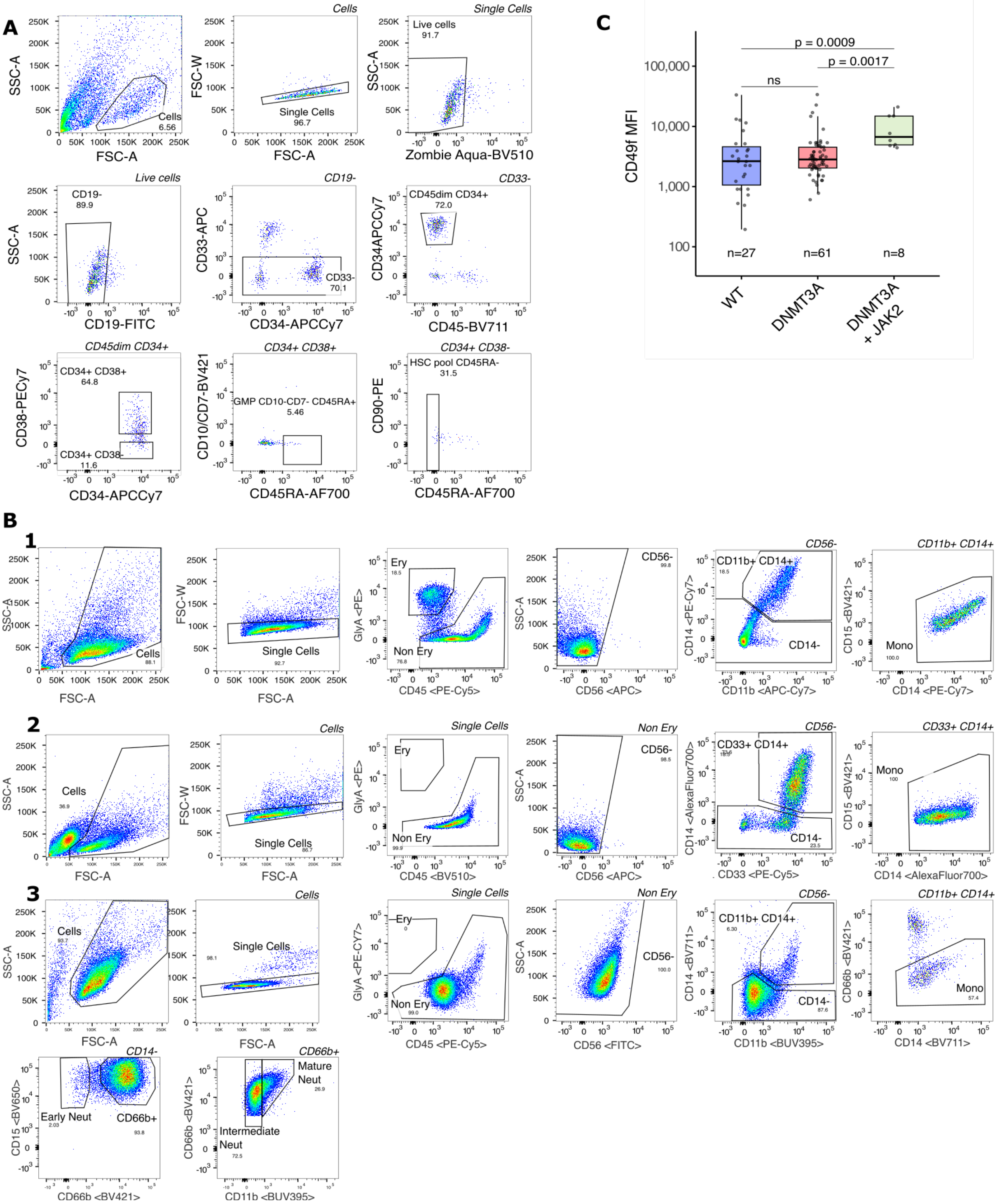


**Figure S1: Flow cytometry panels used in this study and index sorting data from CHIP1. A)** Sorting strategy used to isolate HSPCs from mPB, non-mPB and BM. For AML samples, leukemic blasts were excluded by gating on CD19-/CD33-/CD45dim/CD34+ from viable singlets. HSC/MPPs (CD34+/CD38-/CD45RA-) are cultured *in vitro.* **B)** Gating strategy employed for phenotyping analysis of single-cell derived colonies. The gating strategy was adapted based on the staining panel used (see methods). Representative gating strategy employed for the analysis of **(1)** CHIP1 **(2)** AML1 **(3)** AML2, AML3, AML4, and CHIP5 single-cell derived colonies are shown. **C)** CD49f median fluorescence intensity (MFI) in CHIP1 HSC/MPPs of the indicated genotypes at the time of sort. Genotypes based on the phylogenetic tree shown in **Fig.1B**. mPB: mobilised peripheral blood, BM: bone marrow.

**Figure S2**

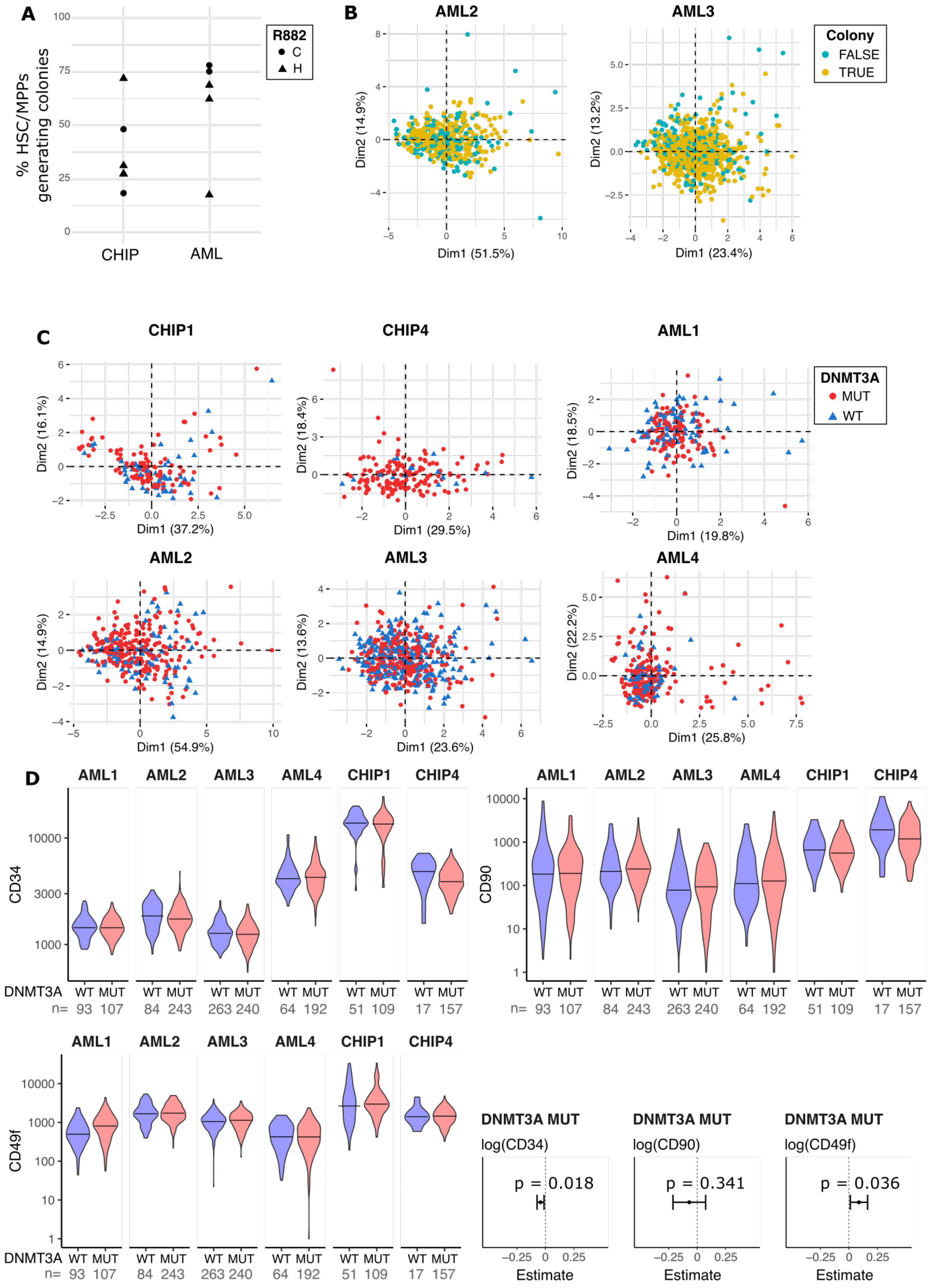


**Fig. S2: Clonogenic efficiency and index sorting analysis of single HSC/MPPs sorted in this study. A)** Clonogenic efficiency (percentage of HSC/MPPs that produced mature colonies amongst all single

HSC/MPPs that were seeded *in vitro)* of all CHIP and AML samples included in this study. **B)** PCA plots derived from 9 cell surface markers on HSC/MPPs indexed at the time of single cell sort. True: cells that produced colonies; False: cells that did not produce a colony. 2 representative examples, AML2 and AML3 shown here. **C)** PCA plots derived from 9 cell surface markers on HSC/MPPs indexed at the time of single cell sort; Blue triangles: WT cells; Red circles: *DNMT3A*-R882 cells. **D**) Violin plots of MFI of cell surface markers of stemness (CD34, CD90 and CD49f) on HSC/MPPs of the indicated genotypes at the time of sorting; number of HSC/MPPs (n) indicated under each violin plot. Forest plot reports estimated *DNMT3A* mutation effect (with 95% confidence interval) on the parameter of interest, obtained by mixed effect modelling (see Methods). p values were computed with Satterthwaite's method and corrected with Benjamini–Hochberg's.

**Figure S3**

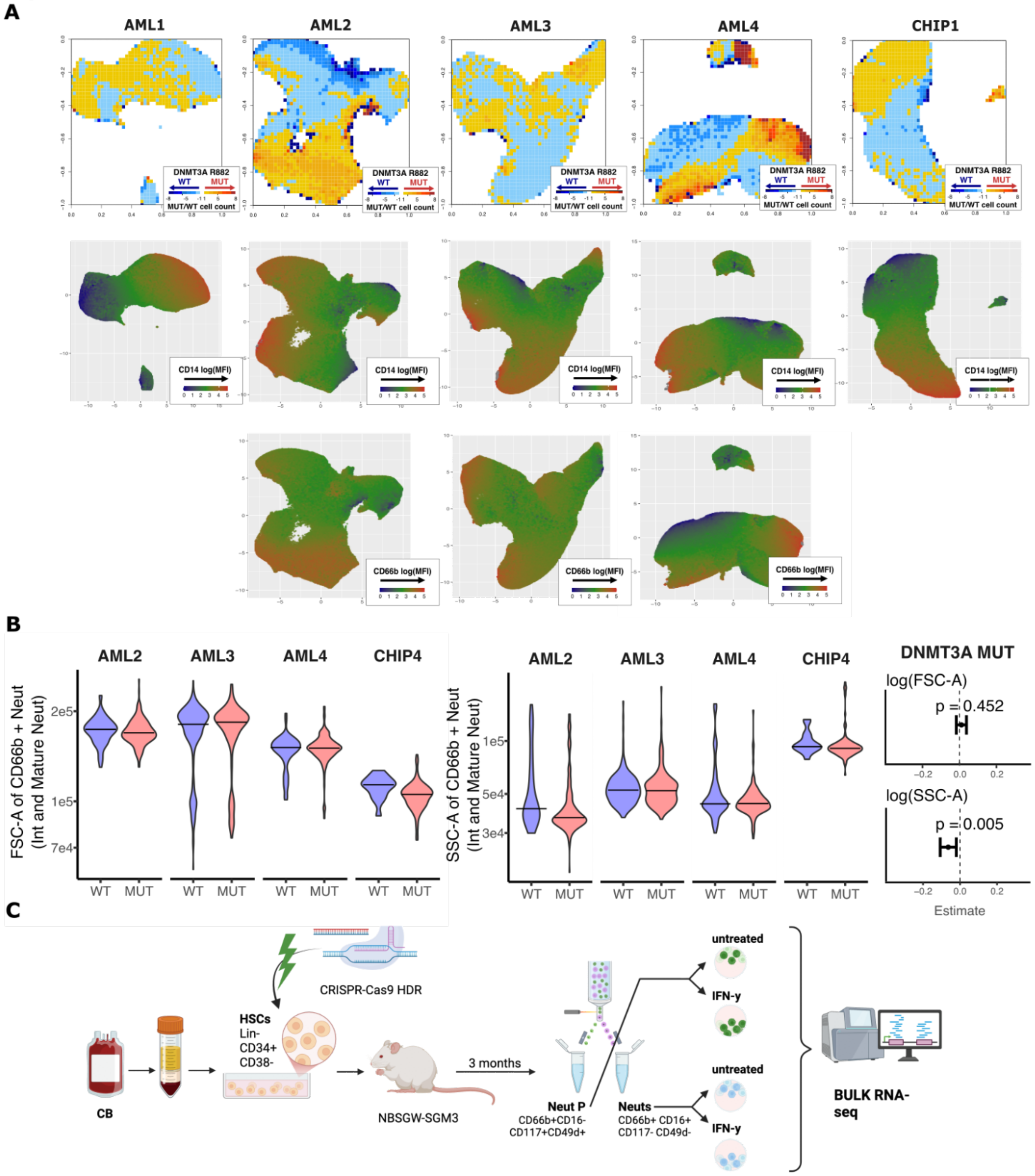


**D**

| MOUSE_ID | %human CD45+ BM cells | GENOTYPE | R882H VAF chr2:25,457,242 | DELETION in chr2: 25,457,247 |
|---|---|---|---|---|
| 80NM | 40.39 | R882H | 37.67 | 1.19 |
| 80R1 | 42.17 | R882H | 42.07 | 2.05 |
| 87R1 | 59.59 | R882H | 16.31 | 20.85 |
| 87R1L1 | 24.73 | R882H | 12.88 | 38.96 |
| W07 | 49.98 | WT | 0.00 | 0.00 |
| W35 | 50.77 | WT | 0.00 | 0.00 |
| W73 | 35.62 | WT | 0.00 | 0.00 |

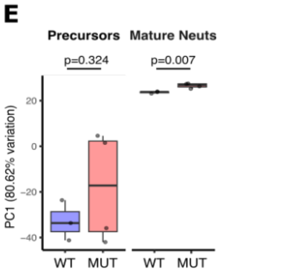


**Fig. S3: Primary sample phenotype analysis and *DNMT3A*-R882H edited humanised mouse model**
**A)** Pooled *in silico* analysis of all mature cells generated *in vitro* from *DNMT3A* MUT and WT HSC/MPPs, per individual. Top panel: ratio of *DNMT3A*-R882 mutated cells / WT cells in each area of the UMAP;

Middle and bottom: MFI of CD66b (middle) and CD14 (bottom) cell surface markers. **B)** Violin plots of physical parameters (FSC-A, SSC-A) of intermediate and mature neutrophils (CD66b+) recorded by flow cytometry analysis. n: number of HSC/MPPs derived colonies containing neutrophils. Forest plot reports estimated *DNMT3A*-R882 mutation effect (with 95% confidence interval) on the parameter of interest, obtained by mixed effect modelling (see Methods). p values were computed with Satterthwaite's method and corrected with Benjamini–Hochberg's. **C)** *DNMT3A*-R882H edited humanised mouse model study design. **D)** Engraftment efficiency and VAF of genomic edits at the *DNMT3A* genomic locus at the time of bone marrow (BM) harvesting **E)** Principal component 1 (PC1) values from transcriptome PCA of untreated neutrophil precursors and mature neutrophils. Welsh's two tailed t-test.

**Figure S4**

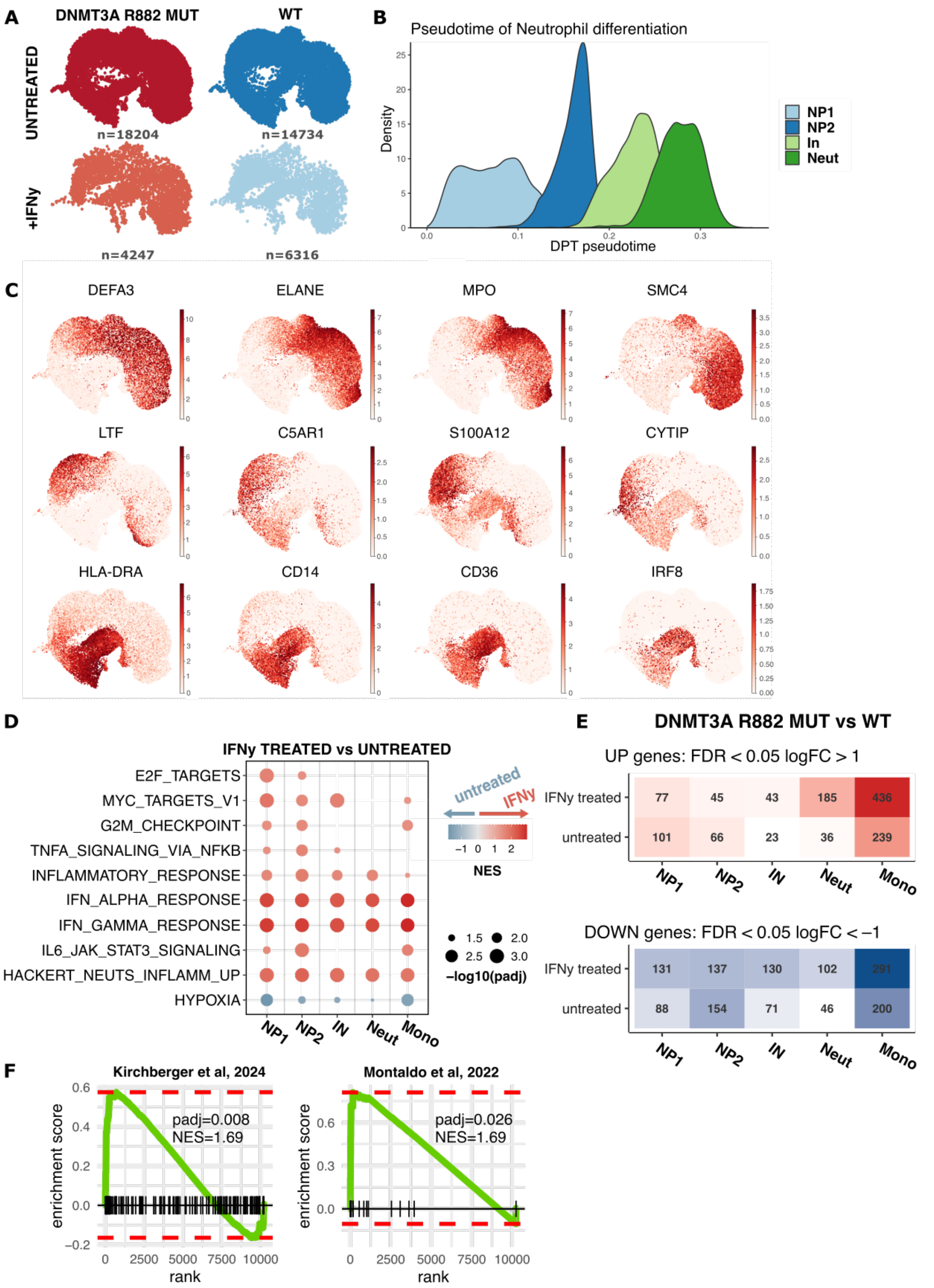


**Fig. S4: scRNA-seq analysis of *in vitro DNMT3A*-R882 and WT haematopoiesis A)** UMAP distribution of cells in every condition. n : number of cells per condition. **B)** Pseudotime analysis of neutrophil lineage clusters. **C)** Representative example of marker genes expression on UMAP **D)** GSEA of Hallmark and selected gene sets (Supplementary Table 3, tab5) in untreated vs IFNγ-treated cells. **E)**

Number of significantly (FDR<0.05) differentially upregulated (logFC>1, top) and downregulated (logFC<-1, bottom) genes per cluster per condition. **F)** GSEA enrichment of neutrophil maturation signatures [53,54] in *DNMT3A*-R882 vs WT IFNγ treated mature neutrophils.
